# Specific excitatory connectivity for feature integration in mouse primary visual cortex

**DOI:** 10.1101/072595

**Authors:** Dylan R Muir, Patricia Molina-Luna, Morgane M Roth, Fritjof Helmchen, Björn M Kampa

**Affiliations:** Biozentrum, University of Basel, Klingelbergstrasse 50/70,4056 Basel, Switzerland.; Laboratory of Neural Circuit Dynamics, Brain Research Institute, University of Zurich, Wjinterthurerstrasse 190, 8057 Zurich, Switzerland.; Department of Neurophysiology, Institute of Biology 2, RWTH Aachen University, Templergraben 55,52062 Aachen, Germany.; JARA-BRAIN, 52074 Aachen, Germany.

**Keywords:** amplification, competition, decorrelation, cortical computation, mouse V1

## Abstract

Local excitatory connections in mouse primary visual cortex (V1) are stronger and more prevalent between neurons that share similar functional response features. However, the details of how functional rules for local connectivity shape neuronal responses in V1 remain unknown. We hypothesised that complex responses to visual stimuli may arise as a consequence of rules for selective excitatory connectivity within the local network in the superficial layers of mouse V1. In mouse V1 many neurons respond to overlapping grating stimuli (plaid stimuli) with highly selective and facilitatory responses, which are not simply predicted by responses to single gratings presented alone. This complexity is surprising, since excitatory neurons in V1 are considered to be mainly tuned to single preferred orientations. Here we examined the consequences for visual processing of two alternative connectivity schemes: in the first case, *local connections are aligned with visual properties inherited from feedforward input* (a ‘like-to-like’ scheme specifically connecting neurons that share similar preferred orientations); in the second case, *local connections group neurons into excitatory subnetworks that combine and amplify multiple feedforward visual properties (a ‘feature binding’ scheme)*. By comparing predictions from large scale computational models with *in vivo* recordings of visual representations in mouse V1, we found that responses to plaid stimuli were best explained by a assuming ‘feature binding’ connectivity. Unlike under the ‘like-to-like’ scheme, selective amplification within feature-binding excitatory subnetworks replicated experimentally observed facilitatory responses to plaid stimuli; explained selective plaid responses not predicted by grating selectivity; and was consistent with broad anatomical selectivity observed in mouse V1. Our results show that visual feature binding can occur through local recurrent mechanisms without requiring feedforward convergence, and that such a mechanism is consistent with visual responses and cortical anatomy in mouse V1.

**Author summary:** The brain is a highly complex structure, with abundant connectivity between nearby neurons in the neocortex, the outermost and evolutionarily most recent part of the brain. Although the network architecture of the neocortex can appear disordered, connections between neurons seem to follow certain rules. These rules most likely determine how information flows through the neural circuits of the brain, but the relationship between particular connectivity rules and the function of the cortical network is not known. We built models of visual cortex in the mouse, assuming distinct rules for connectivity, and examined how the various rules changed the way the models responded to visual stimuli. We also recorded responses to visual stimuli of populations of neurons in anaesthetised mice, and compared these responses with our model predictions. We found that connections in neocortex probably follow a connectivity rule that groups together neurons that differ in simple visual properties, to build more complex representations of visual stimuli. This finding is surprising because primary visual cortex is assumed to support mainly simple visual representations. We show that including specific rules for non-random connectivity in cortical models, and precisely measuring those rules in cortical tissue, is essential to understanding how information is processed by the brain.

## Introduction

Much of our current understanding of local cortical connectivity in neuronal circuits of the neocortex is based on the presumption of randomness. Anatomical methods for estimating connection probabilities [1,2] and techniques for using anatomical reconstructions to build models of cortical circuits [3-7] are largely based on the assumption that connections between nearby neurons are made stochastically in proportion to the overlap between axonal and dendritic arborisations [8].

On the other hand, a wealth of evidence spanning many cortical areas and several species indicates that cortical connectivity is not entirely random. In species that display smooth functional maps in primary visual cortex (V1), such as cat and macaque monkey, long-range intrinsic excitatory connections tend to preferentially connect regions of similar function [9-13]. Although rodents exhibit a mapless, “salt and pepper” representation of basic visual features across V1 [14], nonrandom connectivity is nonetheless prevalent both within and between cortical layers [15-20], reflecting similarities in functional properties [21-25] or projection targets [26-28].

Despite multiple descriptions of specific connectivity in cortex, the rules underlying the configuration of these connections are not entirely clear. Whereas strong connections are more prevalent between neurons with similar receptive fields, the majority of synaptic connections are made between neurons with poorly-correlated receptive fields and poorly correlated responses [24]. This sea of weak synaptic inputs might be responsible for non-feature-specific depolarisation [24] or might permit plasticity of network function [20].

However, another possibility is that weak local recurrent connections reflect higher-order connectivity rules that have not yet been described. Recent reports have highlighted the facilitatory and selective nature of plaid responses in mouse V1 [29-31]. Many neurons in mouse V1 respond to plaid stimuli in accordance with a simple superimposition of their responses to the two underlying grating components (i.e. “component cell” responses; [32]). However, a significant proportion of neurons that are visually responsive, reliable and selective exhibit complex responses to plaid stimuli that are difficult to explain with respect to simple combinations of grating components [30] (Fig. S1).We hypothesised that responses to complex stimuli in mouse V1 could be a result of local combinations of visual features, through structured local recurrent excitatory connectivity. These rules could be difficult to detect through anatomical measurements, if they comprised only small deviations from predominantly “like-to-like” connectivity.

Here we examined whether small tweaks to recurrent connectivity rules could alter visual representations in cortex, by analysing the computational properties of cortical networks with defined rules for local connectivity. We simulated visual responses to grating and plaid stimuli in large networks with properties designed to resemble the superficial layers of mouse V1, assuming distinct connectivity schemes. We then compared the response patterns and visual representations predicted by the network simulations with those recorded *in vivo* in mouse V1, to test the predictions arising from our models.

Specifically, we evaluated two broad classes of connectivity patterns, where specific local excitatory connectivity is defined according to the visual response properties of neurons (Fig. 1):

1. Strictly “like-to-like” connectivity, such that neurons with similar response properties defined by their feed-forward inputs to each neuron (e.g. orientation tuning of neurons in the superficial layers, arising from tuned input from layer 4) are grouped into subnetworks;
2. A form of “feature-binding” connectivity, such that in addition to predominantly “like-to-like” connectivity, excitatory neurons with differing feed-forward visual properties (e.g. distinct orientation preference) are also grouped together.

**Figure 1:**
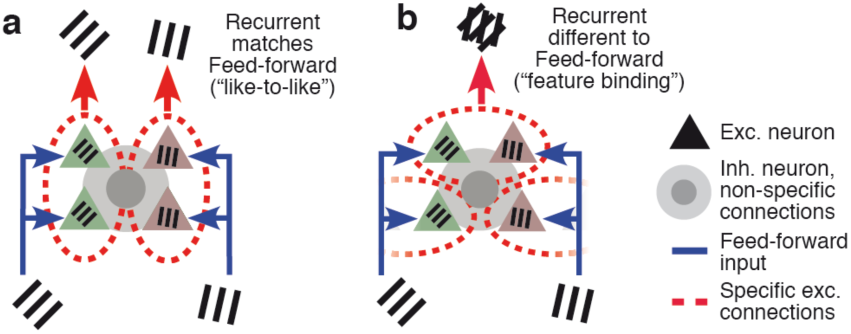
Like-to-like and feature-binding rules for local recurrent connectivity. **a** Connectivity scheme where local recurrent excitatory connections (between neu-rons grouped by dashed ovals) are matched to the feedforward visual preferences of the connected neurons (“like-to-like” over orientation preference, indicated by grating icons). **b** Connectivity scheme where local recurrent excitatory connections are different from the feedforward visual preferences of connected neurons (“feature-binding”). Connections to and from inhibitory neurons (circles and shading) are assumed to be non-specific in all cases. Exe.: excitatory; Inh.: inhibitory.

Despite the small difference in network configuration, these distinct rules give rise to radically different visual representations of plaid stimuli, both in terms of complexity of visual response selectivity of individual neurons and regarding facilitation versus suppression in response to these compound stimuli. We found that the complexity of plaid responses in mouse V1 was reproduced in our simulations when assuming the ‘feature-binding’ connectivity scheme, with local connections grouping multiple feedforward response properties, but not when assuming purely ‘like-to-like’ connections.

## Results

### Models of local connectivity and cortical activity

We designed a non spiking model of the superficial layers of mouse V1, to explore the effect of different connectivity rules on information processing within the cortex. Non-spiking linear-threshold neuron models provide a good approximation to the firing rate / input current (F–I) curves of adapted cortical neurons [33]; model neurons with linear-threshold dynamics can be directly translated into integrate-and-fire models with more complex dynamics [34,35], and in addition form good approximations to conductance-based neuron models [36]. A full list of parameters for all models presented in this paper is given in Table 1.

**Table 1:** Summary of nominal model parameters and model variables. Abbreviations: Exc: Excitatory; Inh: Inhibitory; Prop: proportion.

| Parameter | Description | Nominal value |
| --- | --- | --- |
| $\tau_i$ | Lumped neuron time constant for neuron $i$ | 10 ms |
| $g_j$ | Nominal charge injected by synapses from neuron $j$ | Exc.: 0.01 pC/spike/synapse<br>Inh.: 10×0.01 pC/spike/synapse |
| $\alpha_j$ | Nominal output gain of neuron $j$ | 0.066 spikes/pC |
| $n_{i,j}$ | Number of synapses made from neuron $j$ to neuron $i$ | |
| $\beta_j$ | Threshold of neuron $j$ | Zero |
| $\sigma_i \cdot \zeta_i(t)$ | Noise current injected into neuron $i$. Wiener process with std. dev. $\sigma_i$ after 1 sec. | $\sigma_i = 5$ mA |
| $N_N$ | Number of neurons in simulation | 80,000 (10% of cortical density) |
| Prop. inh. | Proportion of inhibitory neurons | 18% |
|  | Dimensions of simulated torus space | 2.2×2.2 mm |
| $S_i$ | Nominal number of synapses made by neuron $i$ (within superficial layers only) | Exc.: 8142 Inh.: 8566 |
| $\sigma_{d,i}$ | Std. Dev. of Gaussian dendritic field of neuron $i$ | 75μm (approx. width 300μm) |
| $\sigma_{a,i}$ | Std. dev. of Gaussian axonal field of neuron $i$ | Exc.: 290μm (approx. width 1100μm)<br>Inh.: 100μm (approx. width 400μm) |
| $\kappa_i$ | Input orientation tuning width parameter for neuron $i$ | 4 |
| $s_1$ | Degree of like-to-like modulation of anatomical connection probability | |
| $s_2$ | Degree of feature-binding modulation of connection probability | |
| $\kappa_1$ | Orientation tuning of like-to-like connection probability | |
| $\kappa_2$ | Orientation tuning of subnetwork membership probability | |
| $N_s$ | Number of subnetworks that exist at a point in cortex | |
| $\vartheta$ | Number of preferred orientations bound in an subnetwork | |

**Table 2:** Parameter values used to specify large-scale network models.

| Network configuration | Parameter values |
| --- | --- |
| Random connectivity model | $s_1 = 0, s_2 = 0$ |
| Like-to-like specificity model | $s_1 = 0.8, s_2 = 0, \kappa_1 = 0.5$ |
| Feature-binding specificity model | $s_1 = 0.1, s_2 = 0.25, \kappa_1 = 0.5, \kappa_2 = 4, N_s = 6, \vartheta = 2$ |

#### General equations governing model dynamics

Individual excitatory neurons (approximating layer 2/3 pyramidal cells) and inhibitory neurons (approximating layer 2/3 basket cells) were modelled as linear-threshold units, with equal time constants and thresholds set to zero. The dynamics of each rate-coded neuron in the large- and small-scale models was governed by the differential equation

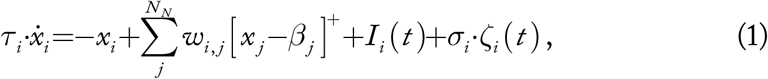

where *τ*_*i*_ is the time constant of neuron *i; x*_*i*_ is the instantaneous current being injected into neuron *i*; [ ]^+^ denotes the linear-threshold transfer function [ *x* ]^+^ = max(*x*, 0); *β*_*j*_ is the activation threshold of neuron *j*; *I*_*i*_ (*t*) is the stimulus input current provided to neuron *i* at time *t*; *σ*_*i*_ *ζ*_*i*_ (*t*) is a white noise process included to approximate the barrage of spontaneous excitatory and inhibitory post-synaptic potentials (EPSPs and IPSPs) experienced by cortical neurons; and *N*_*N*_ is the total number of neurons in the model. The directed connection strength between two neurons *j* and *i* is given in Eq. (1) by *w*_*i*, *j*_ =*g*_*j*_ *n*_*i*, *j*_ · *α*_*j*_, where *g*_*j*_ is the charge injected by a synapse from neuron *j* to neuron *i* and *n*_*i*, *j*_ is the number of synapses made by neuron *j* onto neuron *i*; *α*_*j*_ is the gain of neuron *j*.

#### Synaptic input

Synapses were modelled as constant current sources that injected an amount of charge per second related to the average firing rate of the presynaptic neuron, modulated by the synaptic release probability. Single excitatory synapses were assigned a weight of 0.01 pC / spike / synapse; single inhibitory synapses were considered to be 10 times stronger [4]. Excitatory and inhibitory neurons were assigned output gains of 0.066 spikes / pC [37].

### Specific connectivity gives rise to amplification and competition

The dynamics of neuronal networks defined with particular connectivity rules remain generally unknown, although some results suggest that specific connectivity leads to reduced dimensionality of network activity patterns [38]. Here we explored the relationship between specific connectivity and network dynamical properties in a non-linear, rate-based network model incorporating realistic estimates for recurrent excitatory and inhibitory connection strength in layers 2 / 3 of mouse V1.

To explore the basic stability and computational consequences of functionally specific excitatory connectivity, we built a small five-node model (four excitatory and one inhibitory neurons; “analytical model”; Fig. 2). Connections within this model were defined to approximate the average expected connectivity between populations of neurons in layers 2 / 3 of mouse V1. Excitatory neurons were grouped into two subnetworks, and a proportion *s* of synapses from each excitatory neuron was reserved to be made within the same subnetwork. When *s* = 0, E↔E synapses were considered to be made without specificity, such that each connection in the small model approximated the average total connection strength expected in mouse V1 in the absence of functional specificity. When *s* = 1, all E↔E synapses were considered to be selectively made within the same subnetwork, such that no synapses were made between excitatory neurons in different subnetworks. Connections to and from the inhibitory node were considered to be made without functional specificity in every case, mimicking dense inhibitory connectivity in mouse visual cortex [39-42]. The general form of the weight matrix is therefore given by

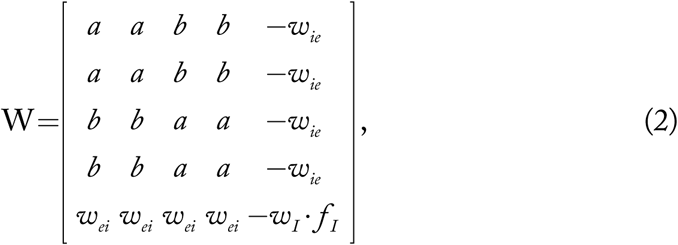

where *w*_*S*_ =*w*_*E*_ · (*1*- *f*_*I*_) *s* is the specific weight component, *w*_*N*_ =*w*_*E*_ · (*1*- *f*_*I*_)·(*1*-*s*) is the nonspecific weight component, *w*_*E*_ is the total synaptic weight from a single excitatory neuron, *w*_*I*_ is the total synaptic weight from a single inhibitory neuron; *f*_*I*_ = 1/5 is the proportion of inhibitory neurons; *a*=*w*_*S*_ / 2+*w*_*N*_ / 4 is the excitatory weight between neurons in the same subnetwork; *b*=*w*_*N*_ / 4 is the excitatory weight between neurons in different subnetworks; *w*_*ie*_ =*w*_*I*_ · (1- *f*_*I*_)/ 4 is the nonspecific inhibitory to excitatory feedback weight; and *w*_*ei*_ =*w*_*E*_ · *f*_*I*_ is the nonspecific excitatory to inhibitory weight.

**Figure 2:**
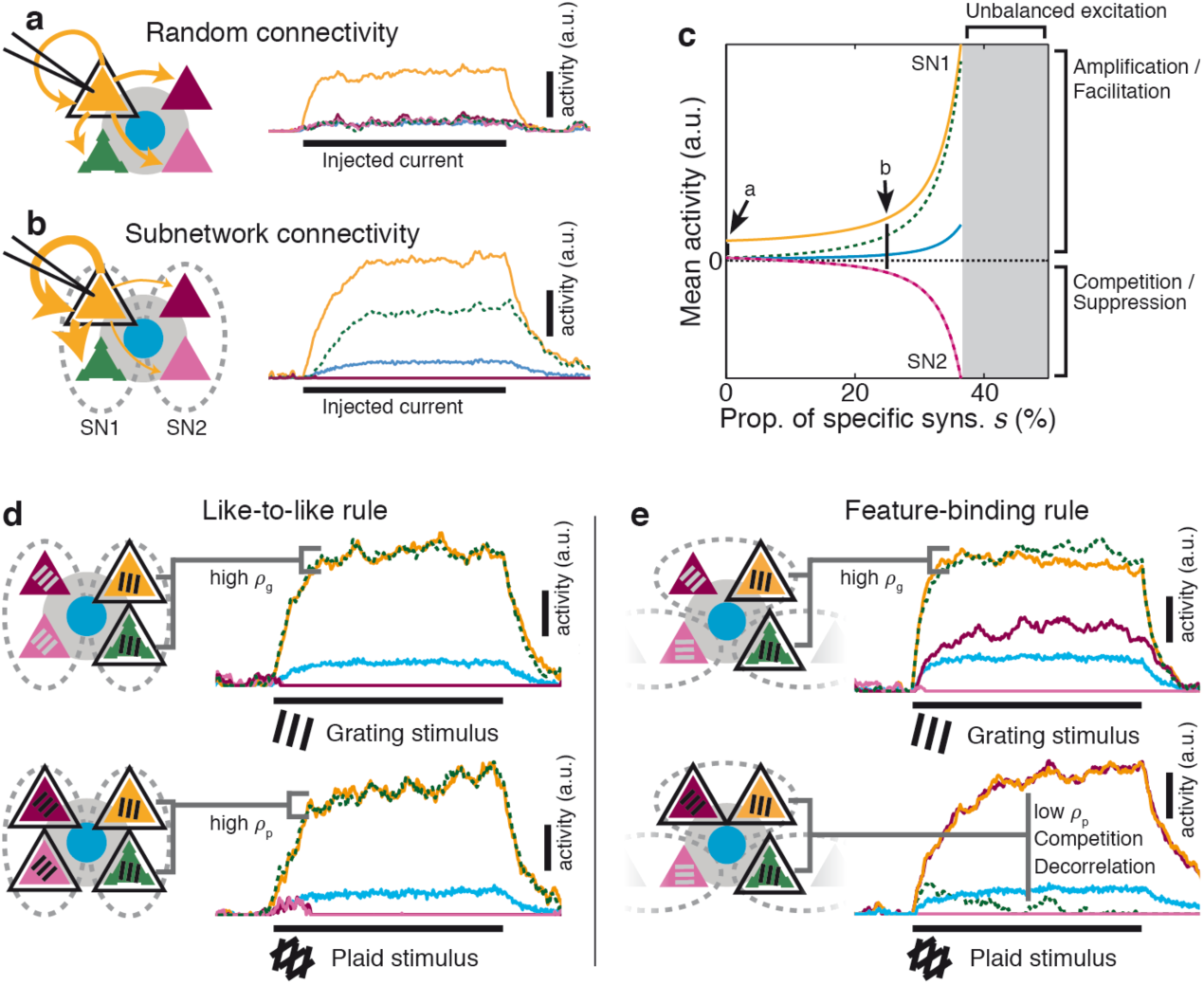
Rules for excitatory connectivity influence stimulus representations, and underlie amplification and competition. **a** In a simple model for random connectivity in mouse Vl, injecting current into a single neuron (black outline) leads to non-specific activation of other excitatory (triangle) and inhibitory neu-rons (circle). Traces show the instantaneous firing rate of each neuron. **b** When the model is partitioned into subnetworks (SNl & 2; dashed ovals), injecting current into a single neuron gives rise to an amplified response within the same subnetwork and suppresses activity in the non-driven subnetwork. **c** The degree of amplification and suppression depends directly on the proportion of excitato-ry synapses *s* restricted to be made within a subnetwork (see Fig. S1). Values of *s* used in panels a-b indicated on plot. d When local recurrent excitatory connections match the feedforward visual properties of connected neurons (“like-to-like”), grating responses (top) and plaid responses (bottom) are highly similar (high *ρ*_*s*_ & *ρ*_*p*_). **e** In contrast, when local recurrent connections are different from the feedforward visual properties-in this case, grouping two different preferred orientations (“feature-binding”)-then neurons with similar grating responses (top, high *ρ*_*g*_) can have dissimilar plaid responses (bottom, low *ρ*_*p*_), reflecting decorrelation of these responses caused by competition. Black outlines: stimulated neurons. Grating labels: preferred orientation of that neuron. Dashed ovals: neurons grouped by specific excitatory connectivity. a.u.: arbitrary units; prop.: proportion; syns.: synapses. Other conventions as in Fig. 1.

#### Measuring stability and competition

To determine network stability in the analytical model, we performed an eigenvalue analysis of the system Jacobian, given by J = (W–I)./T, where W is the system weight matrix as given above, I is the identity matrix, T is the matrix composed of time constants for each post-synaptic neuron corresponding to elements in W and A./B indicates element-wise division between matrices A and B. The network was considered stable if all eigenvalues of J as well as the trace of the Jacobian Tr(J) were non-positive. The non-linear dynamical system was linearized around the fixed point where all neurons are active; if this fixed point is unstable then the system operates in either a hard winner-take-all mode if a different partition is stable, or is globally unstable [43,44]. Neither of these modes is desirable for cortex.

As suggested by estimations of strong excitatory feedback in cortex [4,45], our model required inhibitory feedback to maintain stability (an inhibition-stabilised network or ISN; Fig. S2; [46-49]; but see [50]). For a network to be in an inhibition-stabilised (ISN) regime, the excitatory portion of the network must be unstable in the absence of inhibition, and inhibition must be strong enough in the full network to balance excitation. To determine whether the parameter regimes place the network in an ISN regime, we therefore performed an eigenvalue analysis of the system in which all inhibitory connections were removed (i.e. *w*_*I*_ =0). Either an eigenvalue of the Jacobian J_E_ of the excitatory-only network or the system trace Tr(J_E_) was required to be positive, but the system including inhibitory feedback was required to be stable.

We determined the presence and strength of competition between neurons by injecting current into a single excitatory neuron and recording the net current received by an excitatory neuron in the opposite subnetwork at the network fixed point (see Fig. 2a). Negative net currents correspond to competition between the stimulated and recorded excitatory neurons (shown as shading in Fig. S2). Nonrandom connectivity, in the form of specific excitatory connections within subnetworks (Fig. 2b; SNs; [15,18]), introduced selective amplification within subnetworks and competition between subnetworks (Fig. 2c). Surprisingly, these computational mechanisms were strongly expressed even when only a minority of synapses (*s* around 20%) were made to be subnetwork-specific (Fig. 2c; Fig. S2). Specific connectivity rules resulted in functional grouping of sets of excitatory neurons (Fig. 2b), permitting the network to operate in a soft winner-take-all regime [51,52].

Neither competition nor amplification was present under parameters designed to approximate functionally non-specific connectivity in mouse V1 (Fig. 2a, c; Fig. S2). This is not because the network architecture was incapable of expressing competition, but because recurrent excitatory connections were insufficiently strong under assumptions of random stochastic connectivity. We conclude that specific excitatory connectivity strongly promotes amplification and competition in neuronal responses.

### Local excitatory connections in cortex are broadly selective for preferred orientation

Does the precise configuration of local recurrent connectivity change response patterns in cortical networks? In mouse V1, synaptic connection probability is enhanced by similarity of orientation preference [21,23,25], suggesting that local excitatory connections may group together neurons with common preferred orientations. Connection probability is even more strongly modulated by neuronal response correlations to natural movies; i.e., the likelihood for a synaptic connection is higher for neuronal pairs responding similarly to natural scenes [21,22,24].

If connections in mouse V1 were strictly governed by preferred orientation, then neurons with similar orientation preference should also predominately have similar responses to natural movies, and vice versa. We recorded visual responses populations of neurons labelled with the synthetic calcium indicator OGB in anaesthetized mouse V1 (5 animals, 129 / 391 responsive neurons with overlapping receptive fields / total imaged neurons; Fig. S3a–c; see Methods). We used signal correlations to measure the similarity between the responses of pairs of neurons with identified receptive fields (Fig. S3a) to drifting grating (Fig. S3b) and natural movie (Fig. S3c) visual stimuli (see Methods).

We found that neuronal pairs with high signal correlations to natural scenes, which are most likely to be connected in cortex [21,22,24], showed only a weak tendency to share similar orientation preferences (Fig. S3d–e; pairs with OSI > 0.3; p=0.8, Kruskall-Wallis). This is consistent with earlier findings in cat area 17 (V1), which showed a poor relationship between responses to gratings and natural movies [53].

Similarly, under a like-to-like connectivity rule, synaptically connected neurons in mouse V1 should share both similar orientation preference and responsiveness to natural movies. We therefore compared response correlations and preferred orientations for pairs of mouse V1 neurons, which were known to be connected from *in vivo* / *in vitro* characterisation of functional properties and connectivity (data from [24] used with permission; 17 animals, 203 patched and imaged cells, 75 connected pairs). Consistent with our results comparing responses to gratings and natural movies, connected pairs of cells with similar orientation preference were not more likely to share a high signal correlation to flashed natural scenes (Fig. S3f; p=0.54, Kruskall-Wallis). Also consistent with earlier findings [21,23], we observed a positive relationship between synaptic connectivity and similarity of orientation preference (Fig. S3g; p=0.045, Ansari-Bradley test). However, strongly connected pairs (strongest 50% of excitatory post-synaptic potentials—EPSPs—over connected pairs) were not more similar in their preferred orientation than the remaining pairs (p=0.17, Ansari-Bradley test vs weakest 50% of connected pairs). Connected pairs spanned a wide bandwidth of preferred orientations, with more than 20% of connections formed between neurons with orthogonal preferred orientations. Spatial correlation of receptive fields is a comparatively better predictor for synaptic connectivity than shared orientation preference, but a majority of synaptic inputs are nevertheless formed between neurons with poorly- or un-correlated responses [24]. We conclude that similarity in orientation preference only partially determines connection probability and strength between pairs of neurons in mouse V1.

This weak functional specificity for similar visual properties can be explained by two possible alternative connectivity rules. In the first scenario, local excitatory connections in cortex are aligned with feedforward visual properties, but with broad tuning (Fig. 1a; a “like-to-like” rule). As a consequence, all connections show an identical weak bias to be formed between neurons within similar tuning, and the average functional specificity reported in Fig. S3g and elsewhere [21,24] reflects the true connection rules between any pair of neurons in cortex.

Alternatively, local excitatory connections may be highly selective, but follow rules that are not well described by pairwise similarity in feedforward visual properties. For example, subpopulations of connected excitatory neurons might share a small set of feedforward visual properties, as opposed to only a single feedforward property (Fig. 1b; a “feature-binding” rule). In this case, connections within a subpopulation could still be highly specific, but this specificity would be difficult to detect through purely pairwise measurements. If pairwise measurements were averaged across a large population, any specific tuning shared within groups of neurons would be averaged away.

### Selective amplification under like-to-like and feature-binding connectivity rules

Amplification in the network with specific connectivity is *selective* (Fig. 2b–c): neurons within a subnetwork recurrently support each other’s activity, while neurons in different subnetworks compete. Therefore, which sets of neurons will be amplified or will compete during visual processing will depend strongly on the precise rules used to group neurons into subnetworks. We therefore examined the impact of “like-to-like” and “feature-binding” rules on responses in our analytical model. The excitatory network was partitioned into two subnetworks; connections within a subnetwork corresponded to selective local excitatory connectivity within rodent V1. Under the “like-to-like” rule, neurons with similar orientation preferences were grouped into subnetworks (Fig. 2d).

We tested the response of this network architecture to simulated grating and plaid stimuli, by injecting currents into neurons according to the similarity between the orientation preference of each neuron and the orientation content of a stimulus. Preferred orientations for each excitatory neuron are indicated in Fig. 2. When a stimulus matched the preferred orientation of a neuron, a constant input current was injected (*I*_*i*_ (*t*)=*ι*); when a stimulus did not match the preferred orientation, no input current was provided to that neuron (*I*_*i*_ (*t*)=0). When simulating the analytical model, the input current *ι* =*1*.

Under the “like-to-like” rule, responses of pairs of neurons to simple grating stimuli and more complex plaid stimuli were highly similar (Fig. 2d). Amplification occurred within subnetworks of neurons with the same preferred orientation, and competition between subnetworks with differing preferred orientation [51,54] (visible by complete suppression of response of neurons in lower traces of Fig. 2d).

Alternatively, we configured the network such that the rules for local excitatory connectivity did not align with feedforward visual properties (a “feature-binding” rule). We configured subnetworks by grouping neurons showing preference for either of two specific orientations (Fig. 2e). When this “feature-binding” connectivity rule was applied, neuronal responses to grating and plaid stimuli differed markedly (cf. top vs bottom panels of Fig. 2e). Selective amplification was now arrayed within populations of neurons spanning differing orientation preferences, and competition occurred between subnetworks with different compound feature preferences. Importantly, a “feature-binding” rule implies that neurons with the same preferred orientation could exist in competing subnetworks. While their responses to a simple grating of the preferred orientation would be similar and correlated (Fig. 2e; indicated by a high response correlation measured over grating responses *ρ*_g_), the same two neurons would show decorrelated responses to a plaid stimulus (Fig. 2e; indicated by a low response correlation measured over plaid responses *ρ*_p_). We conclude that changes in pairwise response similarity, provoked by varying the inputs to a network, can provide information about the connectivity rules present in the network.

### Large-scale model of local connectivity in mouse V1

The results of our simulations of the small analytical network suggest that rules for specific local connectivity can modify the correlation of activity between two neurons in a network, depending on the input to the network. The question arises of how connectivity rules shape distributed representations of visual stimuli, examined across a large network and over a broad set of stimuli.

We therefore simulated the presentation of grating and plaid visual stimuli in a large-scale non-linear, rate-based model of the superficial layers of mouse V1. Individual neurons were modelled as described above for the small scale network (Eq. (1)).

To construct the large-scale simulation model of mouse V1, 80,000 linear-threshold neurons were each assigned a random location **u**_*i*_∈𝕋_2_ where 𝕋 defines the surface of a virtual torus of size 2.2 × 2.2 mm. Excitatory and inhibitory neurons were placed with relative densities appropriate for layers 2 and 3 of mouse cortex [55]. Approximately 18% of neurons were inhibitory; [56,57]; see Table 1 for all parameters used in these models. Excitatory neurons were assigned an orientation preference θ drawn from a uniform random distribution, mimicking the “salt and pepper” functional architecture present in rodent visual cortex [14].

#### Anatomical connectivity rules

To determine patterns of synaptic connectivity, we calculated for each neuron the probability distribution of forming a synaptic connection with all other neurons in the model. A fixed number of synapses was drawn from this distribution; the number was chosen as an estimate of the number of synapses formed with other superficial layer neurons in rodent cortex (8142 from each excitatory and 8566 from each inhibitory neuron; [1,55]). Since a simulation with the full density of cortical neurons was computationally infeasible, the size of the simulations was scaled to 10% of estimated cortical density. The sparsity of local synaptic connectivity was maintained by also scaling the number of synapses made by each neuron, while maintaining the total synaptic conductance formed by each neuron.

Axonal and dendritic densities for each neuron were described by a two-dimensional Gaussian field

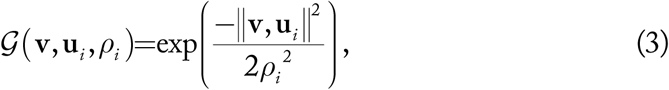

where *ρ*_*i*_ is a field dispersion parameter associated with neuron *i* and ‖**v**, **u‖** is the Euclidean distance between **v** and **u**, computed over the surface of a 2D torus. In our models, each neuron had a Gaussian dendritic field of *ρ*_d_= 75 μm (corresponding to an approximate width of 4ρ = 300 μm; [58]); and axonal field of *ρ*_a,e_= 290 μm for excitatory neurons (width 1100 μm; [58-60]) and *ρ*_a,i_= 100 μm for inhibitory neurons (width 400μm; [61]).

Our default rule for forming synapses was based on Peters’ Rule, in that the probability of forming a synapse was proportional to the overlap between axonal and dendritic fields [2,8]. This was estimated by computing the integrated product of axonal and dendritic fields over a torus 𝕋:

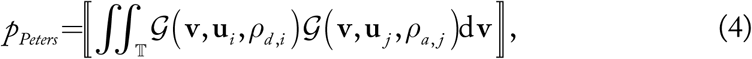

where *p*_*Peters*_ is the probability of forming a single synapse between neurons *i* and *j*, and the notation […] indicates that the expression between the double brackets is normalised to form a probability density function, such that if summed across all possible target neurons the total will be equal to 1.

#### Like-to-like connectivity rule

We investigated two rules for anatomical specificity in intra-cortical excitatory recurrent connections. The first such rule corresponds to the case where local recurrent connectivity is aligned with matching feedforward visual properties (preferred orientation, in our case). We therefore assumed that the probability of forming a synapse is modulated by the similarity in preferred orientation between two excitatory neurons (“Like-to-Like” rule; see Fig. 3a). The probability of connection between two neurons was proportional to

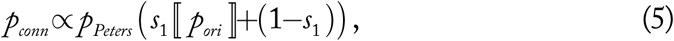

where *p*_*ori*_ =vonmises(*θ*_*i*_, *θ*_*j*_, *κ*); *p*_*Peters*_ is the connection probability under non-specific Peters’ rule connectivity, defined above; and *s*_1_ is the proportional strength of specificity *s*_1_∈[ 0,1]. If *s*_1_ = 0 then Eq. (5) becomes equivalent to Peters’ rule. When *s*_1_ = 1 then the probability of connecting orthogonally tuned neurons is zero.

**Figure 3:**
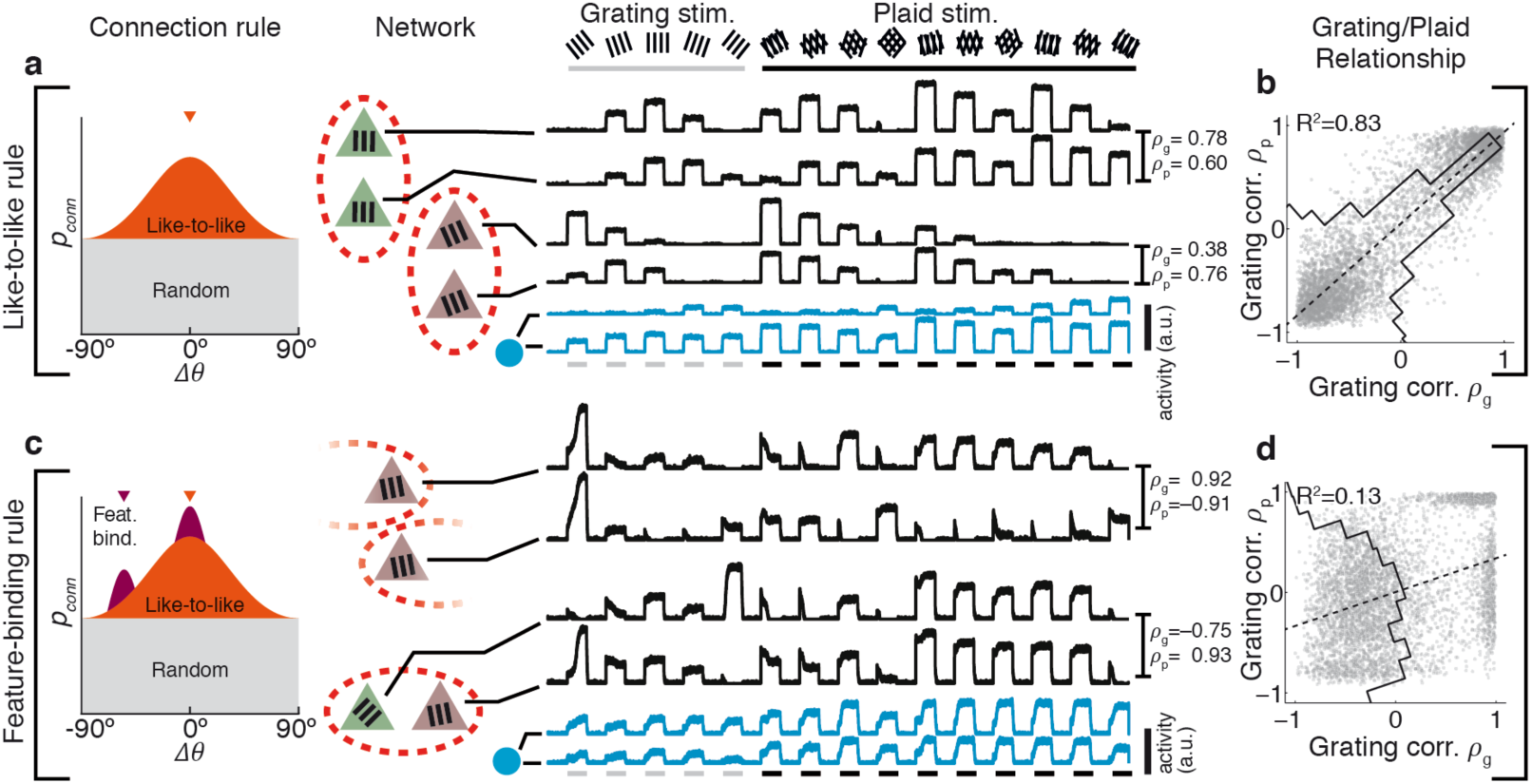
Rules for excitatory connectivity determine response correlation and decorrela-tion in a model of mouse VI. **a-b** In a large-scale network simulation incorporating like–to-like selective excitatory connectivity (connectivity rule and network schematic shown at left), responses of pairs of neurons to grating and plaid stimuli are always similar (b; similar *ρ*_*g*_ & *ρ*_*p*_, high R^2^). Traces: instantaneous firing rates for single example excitatory (black) and inhibitory (blue) neurons. Responses to grating stimuli are highly predictive of plaid responses; distribution of *ρ*_*g*_ versus *ρ*_*p*_ is therefore clustered around the diagonal (black line in b; high R^2^). **c-d** When in addition to like-to-like connectivity, subnetworks also group neurons with several preferred orientations, then pairs of neurons with similar preferred orientations can respond differently to plaid stimuli, and vice versa (see response traces). **d** Competition due to feature-binding connectivity leads to decorrelation of the population response (low R^2^). The distribution of *ρ*_*g*_ versus *ρ*_*p*_ is broad (black line in d), indicating poor predictability between grating and plaid responses. Inhibitory responses are broadly tuned in both models (blue traces in a & b). Pips in connectivity diagram in c indicate example preferred orientations of a single subnetwork. Conventions as in Fig. 1. Stirn.: stimuli; a.u.: arbitrary units; corr.: correlation; feat. bind.: feature binding.

#### Feature-binding connectivity rule

The second rule for anatomical connection specificity corresponds to the case where local recurrent connectivity is not aligned with feedforward visual properties. Instead, it was designed to explore binding of simple visual features (“Feature-Binding” specificity; see Fig. 3e). Under this rule, a subnetwork combined neurons with a number *ϑ* of different orientation preferences. The preferred orientations used to compose a subnetwork in the Feature-Binding specificity model were chosen from periodic filtered noise fields.

Each noise field **Z**_*k*, *q*_ was built by generating a unit-magnitude complex number **z**_*j*_ =exp(-i*ζ*_*j*_) for each neuron in the model, with uniformly-distributed orientations *ζ*_*j*_ ∈[-*π*, *π*). Here “i” represents the complex number 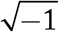; *k*∈[*1, N*_*S*_ ], where *N*_*S*_ is the number of subnetworks in the model; *q*∈[*1,ϑ*], where *ϑ* is the number of preferred orientations per subnetworks. In our models described in this paper, *N*_*S*_ =6 and *ϑ* =2.

A field **Z**_*k*, *q*_ was defined by placing each **z**_*j*_ at the location **u**_*j*_ of the corresponding neuron. Each complex field **Z**_*k*, *q*_ was spatially filtered by convolving with a Gaussian field G_*ρ*_ on a torus, with a spatial standard deviation of *ρ* =75 μm (width 300 μm). The angles from the resulting field of complex numbers was used as one orientation component for one subnetwork, at each point in simulated space. The composition of each subnetwork therefore changed smoothly across cortical space, so that nearby neurons in the same subnetwork had similar functional selectivity. Therefore, ∠(**Z**oG_*ρ*_) defines a *N*_*S*_×*ϑ* matrix of numbers where each element determines one preferred orientation component of the corresponding subnetwork.

Neurons were assigned to one of *N*_*S*_ subnetworks, according to the maximum similarity between a neuron’s preferred orientation and the orientation composition of the set of subnetworks at the location of the neuron’s soma. The similarity between a neuron’s preferred orientation and a subnetwork orientation was computed using a von Mises function (a circular, Gaussian-like function) with width parameter *κ*_2_, such that the membership probability was proportional to

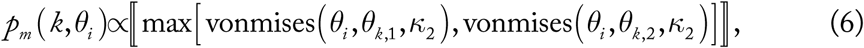

where 𝓀 is the index of an SSN consisting of preferred orientations *θ*_*k*,1_ and *θ*_*k*, 2_; *θ*_*i*_ is the preferred orientation of a neuron under consideration; and the expression within the double brackets ⟦…⟧ was normalised to be a valid probability density function over k. A neuron was assigned membership of an SSN according to the formula

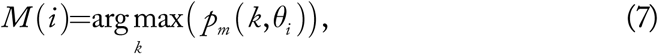

where *M* (*i*) gives the index of the SSN of which neuron i is a member.

The probability of connection between two neurons under the feature-binding model is therefore given by

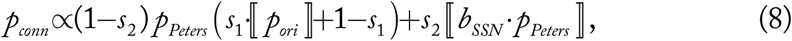

where parameter *s*_1_ determines the relative contribution of Non-Specific versus orientation-tuned Like-to-Like specificity as in Eq. (5); *s*_2_ determines the relative contribution of Feature-Binding specificity; *p*_*ori*_ =vonmises(*θ*_*i*_, *θ*_*j*_, κ_1_) as in Eq. (5); and b_SSN_ is a value equal to 1 when the two neurons fall within the same subnetwork; that is

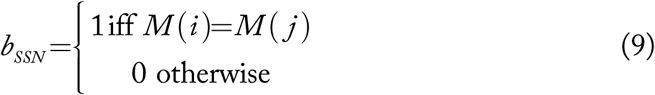

#### Network input

Input was provided to the network as a simulation of orientation-tuned projections from layer 4 to layers 2/3 [62,63]. Each excitatory neuron was assigned an orientation tuning curve based on a von Mises function, with a randomly chosen preferred orientation *θ*_*i*_ and a common input tuning curve width κ = 4. vonmises(·) is the non-normalised von Mises function with vonmises(·)∈[ 0,1], given by

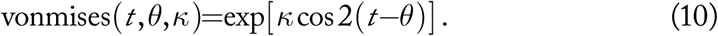

Current was injected into each simulated neuron proportional to the orientation tuning curve of that neuron, according to the orientation content of the stimulus:

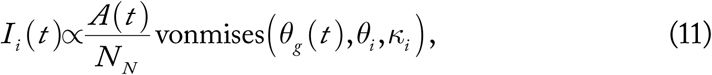

where *A*(*t*) is the amplitude of the stimulus at time *t*; *θ*_*g*_ (*t*) is the orientation of a grating stimulus at time *t*; *θ*_*i*_ is the preferred orientation of neuron *i*; κ_*i*_ is the tuning curve width of neuron *i*; *N*_*N*_ is the total number of neurons in the network. The input to the network is normalised such that the total current injected into the network is equal to *A*(*t*). For a simulated plaid stimulus composed of the two component orientations 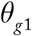 and 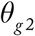, input to a neuron was the linear average of input associated with each grating component, given by

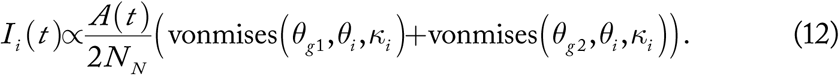

Both grating and plaid stimuli were considered to cover the full visual field. Tuned input currents were injected only into excitatory neurons, because we wanted to investigate the effect of excitatory recurrence on cortical information processing. Providing untuned feedforward input to inhibitory neurons can produce the illusion of competition between excitatory neurons, merely due to the thresholding effect of feedforward inhibitory input shared between those neurons.

#### Inclusion of experimental response variability

We simulated large-scale networks as described above, and obtained responses to simulated visual stimuli. In order to mimic the response variability due to experimental conditions, such as recording noise and intrinsic neuronal response variability, we introduced a random component to the model responses.

To quantify experimental variability, we recorded neuronal responses to presented visual stimuli under two-photon calcium imaging in mouse V1 (see Methods). For each presented stimulus *i* (e.g. a grating of a given orientation), we obtained a set *S*_*i*_ of single-trial responses *r*_*i,j*_ for a single neuron such that *r*_*i*, *j*_ ∈*S*_*i*_, and the trial-averaged response 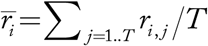, where *T* is the number of trials collected for that stimulus. Over the full set of stimuli for a given neuron, we determined the maximum trial-averaged response 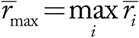. We then measured the standard deviation *σ* over the collection of all single-trial responses over all stimuli for a given neuron normalised by 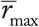, such that 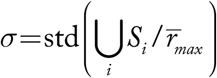. The estimated experimental variability 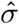 was defined as the median *σ* over all recorded neurons.

A similar procedure in reverse was applied to model-simulated visual responses, to mimic experimental variability. Activity of single neurons in response to a simulated stimulus i was interpreted as the mean response 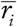, with 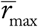 defined as above. Single-trial model responses were then generated as 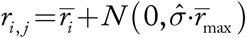, where *N* (*μ*,*σ*) generates a single normally-distributed random variate with mean *μ* and standard deviation *σ*. Twelve trials were generated for each stimulus (i.e. *T*=12), and single-trial responses were then analysed as described for experimentally recorded responses.

#### Estimation of parameters for connection rules

Ko and colleagues characterised functional specificity in mouse V1, by recording in slice from pairs of neurons that were functionally characterised *in vivo* [21]. We fit our function *p*_conn_ (Eq. (5)) to their measurements of the probability of connection between neurons tuned for orientation, giving estimates for both *k*_1_ and *s*_1_ (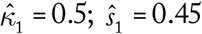). These parameters correspond to fairly weak functional specificity. We found that in the Like-to-Like specificity model, in order to have an appreciable network effect we had to increase the strength of functional specificity to *s*_1_ = 0.8 (with κ_1_ = 0.5). The connectivity measurements of Yoshimura and colleagues suggest that on the order of *N* = 5–6 subnetworks exist in layers 2/3 of rodent cortex [15]. For the Feature-Binding specificity model, we took the parameters *s*_1_ = 0.45, *s*_2_ = 0.225, κ_1_ = 0.5, κ_2_ = 4, *N* = 6, ϑ = 2.

Therefore, in our “feature-binding” model, the majority (68%) of recurrent synapses are made randomly; a smaller fraction (27%) are made according to similarity of preferred orientation and the remaining small fraction (5%) are made selectively across preferred orientations. These last few synapses could potentially weakly change the preferred orientation of a neuron. However, we found that most neurons in our feature-binding model had grating responses aligned with their feed-forward preferred orientation. This is likely due to the strong influence of like-to-like connectivity even in the feature-binding model.

### Feature-binding connectivity leads to facilitation and decorrelation in large networks

We simulated the presentation of grating and plaid stimuli in our large-scale network model of mouse V1. We quantified response similarity between pairs of neurons as suggested by the results of the small network simulations: by measuring pairwise response correlations over a set of grating stimuli (*ρ*_g_), and separately over a set of plaid stimuli (*ρ*_p_; see Methods).

In the network that implemented a “like-to-like” connection rule for recurrent excitatory connectivity (Fig. 3a–b), pairs of neurons showed similar responses to both grating and plaid stimuli (Fig. 3b; R_2_=0.83 between *ρ*_g_ and *ρ*_p_), in agreement with the analytical “like-to-like” model of Fig. 2d.

However, in the network that implemented a “feature-binding” connection rule, where in addition to spatial proximity and similarity in preferred orientation subnetworks were defined to group neurons of two distinct preferred orientations (Fig. 3c–d), neurons showed reduced correlation in response to plaid stimuli (Fig. 3d, R_2_=0.13 between *ρ*_g_ and *ρ*_p_), in agreement with the analytical “feature-binding” model of Fig. 2e. Different configurations of local recurrent excitatory connectivity produced by “like-to-like” or “feature-binding” rules can therefore be detected in large networks, by comparing responses to simple and compound stimuli.

Consistent with our analytical models, networks without functionally specific connectivity did not give rise to decorrelation (Fig. S4b; R^2^=0.72 between *ρ*_g_ and *ρ*_p_). This shows that decorrelation between plaid and grating responses in our models does not arise simply due to random connectivity, but requires the active mechanism of selective amplification through feature-binding subnetwork connectivity.

Inhibitory responses were untuned in our simulations (blue traces in Fig. 3a, c), in agreement with experimental observations of poorly-tuned inhibition in mouse V1 [40,57,64,65].

### Visual responses in mouse V1 are consistent with “feature-binding” connection rules

Our analytical network results show that in principle the configuration of local excitatory connectivity, whether aligned with or spanning across feedforward visual properties, has a strong effect on visual representations (Fig. 2). Our large-scale simulations show that these effects can be detected in large networks as differences in the pairwise correlations of responses to simple and compound visual stimuli (Fig. 3). We therefore aimed to test which connectivity scheme is more likely to be present in visual cortex, by examining responses of neurons in mouse V1.

Using two-photon calcium imaging, we recorded responses of populations of OGB-labelled neurons in mouse V1 to a set of contrast-oscillating oriented grating stimuli over a range of orientations, as well as the responses to the set of plaid stimuli composed of every possible pair-wise combination of the oriented grating stimuli (Fig. 4a; 5 animals, 5 sessions, 313 / 543 responsive / total imaged neurons; see Methods). Responses to plaid stimuli in mouse V1 suggest that stimulating with a denser sampling of compound stimulus space leads to a better characterisation of response selectivity [31] (Fig. S1). Accordingly, we probed responses in mouse V1 under stimuli analogous to those used in the model simulations, with a dense coverage of plaid combinations over a set of finely-varying grating orientations.

**Figure 4:**
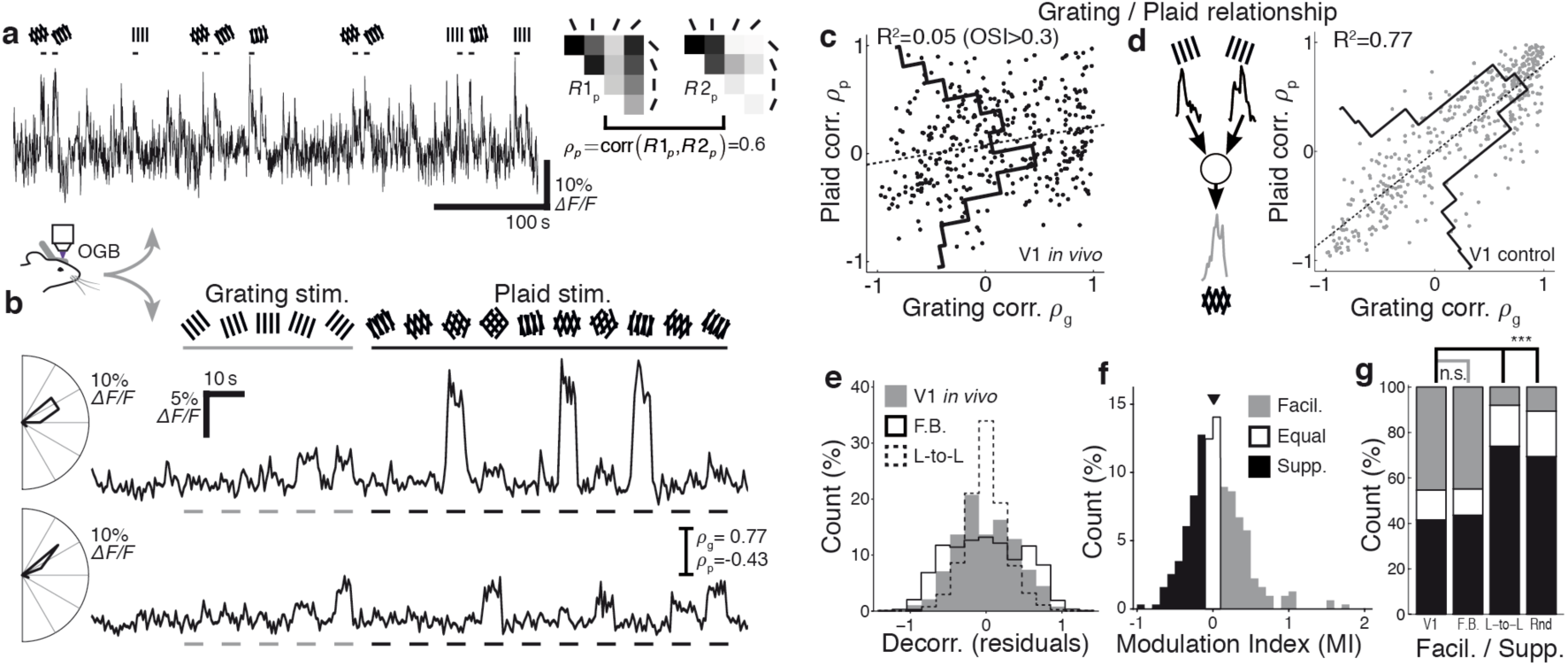
Responses to contrast-oscillating plaid and grating stimuli in mouse VI suggest feature-binding connection rules. **a** Single-trial 0GB calcium response to contrast-oscillat-ing grating and plaid stimuli; presentation time of stimuli evoking strong responses indicated above trace. Right inset: measurement of plaid response similarity ρ between two neurons. **b** Trial-averaged responses (8 trials) of a pair of neurons from a single imaging site, with similar preferred orientations (polar plots at left; high *ρ*_g_) but with dissimilar responses to plaid stimuli (low *ρ*_P_). **c** Responses to grating and plaid stimuli are poorly related in orientation-tuned neurons in mouse Vl (Broad distribution of *ρ*_g_ versus *ρ*_p_ residualsblack line, low R^2^). **d** Control data that includes experimental noise and response variability, obtained by resampling experimental responses and assuming a like-to-like connectivity rule (inset; see Methods), predicts a strong relationship between grating and plaid representations (high R^2^) and is easily distinguished from observed Vl responses in c. **e** Decorrelation in mouse Vl is similar to the "feature-binding” model (F.B.), and much broader than the "like-to-like” mod-el (L-to-L). **f** Responses to plaid stimuli in Vl are split between facilitating and suppressing (45% MI> 0.05; 42% MI< −0.05). **g** The distribution of facilitating (Facil.; MI> 0.05) and sup-pressing (Supp., MI< −0.05) responses is similar between mouse Vl and the "feature-binding” model (F.B.; p = 0.17, Fisher’s exact test). The "like-to-like” and random non-specific (Rnd) connectivity models produced predominately suppressing responses. ***p< 0.001. nVl = 313; nF.B. = 804; nL-to-L = 729; nRnd = 729; significantly responsive neurons with OSI> 0.3. Stirn: stimuli; corr.: correlation; decorr.: decorrelation.

We found that consistent with our earlier findings examining 90° drifting plaid stimuli [30], responses to grating stimuli did not well predict responses to plaid stimuli. Pairs of neurons with similar preferred orientation but with highly differing responses to plaid stimuli were common (Fig. 4b–c; R^2^=0.05 between *ρ*_g_ and *ρ*_p_; OSI > 0.3). The degree of decorrelation we observed in mouse V1 was considerably higher than predicted by the “like-to-like” model, and was more consistent with the “feature-binding” model (Fig. 4e).

Decorrelation induced by plaid responses and the lack of a relationship between grating and plaid responses in mouse V1 were not a result of unreliable or noisy responses *in vivo*. We included in our analysis only neurons that were highly reliable, and responded significantly more strongly than the surrounding neuropil (see Methods). As a further control, we used experimentally recorded responses to grating stimuli to generate synthetic plaid responses for mouse V1 that would result from a cortex with like-to-like subnetwork connectivity (Fig. 4d, inset; see Methods). Our control data were generated from single-trial responses of single V1 neurons, and therefore included the same trial-to-trial variability exhibited by cortex. This control analysis indicates that a “like-to-like” rule among V1 neurons would result in a higher correlation of grating and plaid responses than experimentally observed (Fig. 4d; median R^2^= 0.77 ± [0.767 0.775] between *ρ*_g_ and *ρ*_p_; n = 2000 bootstrap samples; compared with R^2^= 0.05 for experimental results; p < 0.005, Monte-Carlo test).

Importantly, this control analysis is not restricted to our “like-to-like” rule, but makes similar predictions of highly correlated grating and plaid responses for any arbitrary model that combines grating components to produce a plaid response, as long as that rule is identical for every neuron in the network [30]. This is because if a single consistently-applied rule exists, then any pair of neurons with similar grating responses (high *ρ*_g_) will also exhibit similar plaid responses (high *ρ*_p_). In contrast, neurons that are connected within the “feature-binding” model combine different sets of grating components, depending on which subnetwork the neurons are members of.

Neurons in mouse V1 exhibited a wide range of facilitatory and suppressive responses to plaid stimuli, roughly equally split between facilitation and suppression (Fig. 4f–g; 45% vs 42%; MI > 0.05 and MI < –0.05). The proportion of facilitating and suppressing neurons in mouse V1 was similar to that exhibited by responsive neurons in our “feature-binding” model (Fig. 4g; V1 versus F.B., p = 0.17; two-tailed Fisher’s exact test, n_V1_ = 313, n_F.B._= 809). In contrast, neither the “like-to-like” model (L-to-L) nor a model without functionally specific connectivity (Rnd) exhibited significant facilitation in responsive neurons, and both were significantly different from the distribution of facilitation and suppression in mouse V1 (Fig. 4g; p < 0.001 in both cases; two-tailed Fisher’s exact test, n_L-to-L_= 729, n_Rnd_= 729). The wide range of facilitatory and suppressive responses observed in mouse V1 is more consistent with a feature-binding rule for local connectivity, compared with a like-to-like rule or a network without functionally specific connectivity.

## Discussion

Whereas feedforward mechanisms for building response properties in visual networks have been extensively studied, it is not well understood how visual responses are shaped by local recurrent connections. We hypothesised that the configuration of local recurrent cortical connectivity shapes responses to visual stimuli in mouse V1, and examined two alternative scenarios for local connection rules: essentially, whether local excitatory connections are made in accordance with feedforward visual properties (“like-to-like”; Fig. 1a), or whether local excitatory connections span across feedforward visual properties to group them (“feature-binding”; Fig. 1b). We found that highly selective and facilitatory responses to plaid stimuli observed in mouse V1 (Fig. S1, Fig. 4; [30]) are consistent with tuning of recurrent connections within small cohorts of neurons to particular combinations of preferred orientations. Moreover, responses in mouse V1 are inconsistent with a simple configuration of cortical connections strictly aligned with feedforward visual responses.

### Detecting feature-binding connectivity rules in cortex

We found that the precise rules that determine local connections among neurons in cortex can strongly affect the representation of visual stimuli. The “feature-binding” rule we examined embodies the simplest second-order relationship between connectivity and preferred orientation, and was chosen for this reason. We cannot rule out more complicated connectivity rules as being present in mouse V1, but we have shown that a simple “like-to-like” rule cannot explain responses to plaid visual stimuli. Random, non-functionally specific connections were also unable to explain complex plaid responses in mouse V1 (Fig. S4).

How can the detailed statistics of “feature-binding” rules be measured in cortex? Existing experimental techniques have been used to measure only first-order statistical relationships between function and cortical connectivity [18,21-24,40]. Unfortunately, current technical limitations make it difficult to measure more complex statistical structures such as present under a “feature-binding” connectivity rule. Simultaneous whole-cell recordings are typically possible from only a small numbers of neurons, thus sparsely testing connectivity within a small cohort. Even if simultaneous recordings of up to 12 neurons are used [17], identifying and quantifying higher-order statistics in the local connectivity pattern is limited by the low probability of finding connected excitatory neurons in cortex. Nevertheless, our “feature-binding” connectivity model is consistent with the results of functional connectivity studies (Fig. S3).

In addition, our results highlight that small changes in the statistics of local connectivity can have drastic effects on computation and visual coding. Introducing a small degree of specificity, such that a minority of synapses are made within an excitatory subnetwork, is sufficient to induce strong specific amplification and strong competition to the network, even though a majority of the synapses are made randomly without functional specificity (Fig. 2a–c). Under our “feature-binding” model 68% of synapses are made randomly; approximately 27% are made under a “like-to-like” rule and the remaining 5% are used to bind visual features.

Clearly, detecting the small proportion of synapses required to implement feature binding in V1 will be difficult, using anatomical sampling techniques that examine only small cohorts of connected neurons.

A recent study functionally characterised the presynaptic inputs to single superficial-layer neurons in mouse V1, using a novel pre-synaptic labelling technique [66]. Consistent with our results for preferred orientation (Fig. S3f, g), they found that presynaptic inputs were similarly tuned as target neurons but over a wide bandwidth. The majority of synaptically connected networks were tuned for multiple orientation preferences across cortical layers, similar to the feature-binding networks in our study.

We implemented an alternative approach, by inferring the presence of higherorder connectivity statistics from population responses in cortex. This technique could be expanded experimentally, by presenting a parameterised battery of simple and complex stimuli. Stimuli close to the configuration of local connectivity rules would lead to maximal facilitation and competition within the cortical network. Importantly, our results strongly suggest that simple stimuli alone are insufficient to accurately characterise neuronal response properties in visual cortex.

### Amplification and competition might underlie facilitation and suppression

Our theoretical analysis and simulation results demonstrate that functionally specific excitatory connectivity affects the computational properties of a cortical network by introducing amplification of responses within subnetworks of excitatory neurons, and competition in responses between subnetworks (Fig. 2a–c). Several recent studies have demonstrated that visual input is amplified within the superficial layers of cortex [67-69], and recent results from motor cortex suggest competition between ensembles of neurons [70]. Our modelling results indicated that some form of selective local excitatory connectivity is required for such amplification to occur through recurrent network interactions, with reasonable assumptions for anatomical and physiological parameters for rodent cortex (Fig. 2a–c; Fig. S2). This still leaves in question whether the *particular configuration* of selective excitatory connectivity plays a role.

Our simulation results showed that the effects of amplification and competition on cortical responses are tuned to the statistics of local connectivity. This implies that complex visual stimuli for which the composition of stimulus components matches the statistics of a subnetwork will undergo stronger amplification than other non-matching visual stimuli (Fig. 5). In our “feature-binding” model, the statistics of subnetwork connectivity were defined to reflect combinations of two preferred orientations chosen from a uniform random distribution. This combination of two orientations is similar to the visual statistics of plaid stimuli with arbitrarily chosen grating components. As a result, plaid stimuli gave rise to stronger amplification than single grating components alone, when the composition of the plaid matched the composition of connectivity within a particular subnetwork. This led to a facilitatory effect, where some neurons responded more strongly to plaid stimuli than to the grating components underlying the plaid stimuli. Conversely, competition between subnetworks led to weaker responses to some plaid stimuli, for neurons that “lost” the competition. Competition could therefore be one cortical mechanism underlying cross-orientation suppression in response to plaid stimulation.

**Figure 5:**
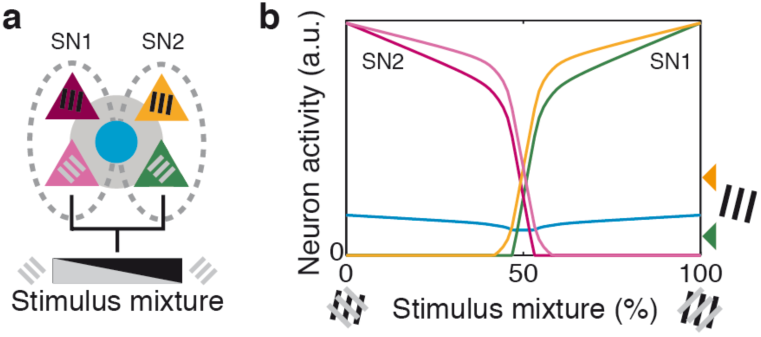
Non-random connectivity supports autoassociative behaviour. In a simple model with two subnetworks **(a),** presenting a linear graduated mixture between the ideal stimuli for the two subnetworks **(b)** results in competition and switching between network representations. When the stimulus is ideal for one subnetwork (mixture= 0% or 100%), then strong amplification ofthe network response occurs (compare with response ofSN 1 to a single grating component; arrowheads at right ofb). When an approximately even mixture is presented (above and below 50%), the network switches rapidly from one representation to the other. Proportion ofspecific excitatory synapses *s*= 25%. Dashed ovals: neu-rons grouped by specific excitatory connectivity. Other conventions as in Fig. 1. a.u.: arbitrary units.

In contrast, suppression in the “like-to-like” and “random non-specific” models occur because the energy in the stimulus is spread across two grating components, and is not combined by the network to form strong plaid selectivity. In the “liketo-like” model, competition occurs between representations of the two oriented grating components of the plaid, causing additional suppression. The presence of amplified, strongly facilitating plaid responses in mouse V1 is therefore consistent with the existence of subnetworks representing the conjunction of differently-oriented edges.

### Building plaid responses from convergence of simple feedforward inputs, or from feedback inputs

Could the complexity of plaid texture responses in mouse V1 be explained by convergence of differently tuned feedforward inputs from layer 4 onto single layer 2/3 neurons, similar to the proposed generation of pattern-selective responses in primate MT [32,71]? Building plaid responses in this way would imply that layer 2/3 neurons would respond to multiple grating orientations, since they would receive approximately equal inputs from at least two oriented components. However, layer 4 and layer 2/3 neurons are similarly tuned to orientation in rodent V1 [62,63], in conflict with this feedforward hypothesis.

In addition, if responses to complex stimuli were built by feedforward combination of simple grating components, then the response of a neuron to the set of grating stimuli would directly predict the plaid response of that neuron. This would then imply that two neurons with similar responses to plaid stimuli must have similar responses to grating stimuli. However we found this not to be the case experimentally; two neurons with similar responses to grating components often respond differently to plaid textures or to natural scenes (Fig. S3d; Fig. 4a, b;[30]).

We cannot rule out the influence of feedback projections on shaping responses to plaid stimuli. The time resolution of calcium imaging is too slow to differentiate between feedforward, recurrent local, and feedback responses based only on timing. However, top-down feedback inputs are considered to be suppressed during anaesthesia [72]; in contrast, we observed complex responses to plaid stimuli in anaesthetised animals. Since our proposed mechanism for feature binding relies on recurrent amplification, relatively few excitatory synapses are required to reproduce complex plaid responses. In contrast, non recurrent influences such as feedforward or feedback projections would require comparatively more synapses to achieve a similar pattern of plaid responses. There are more local recurrent excitatory synapses in V1 layer 2 / 3 than there are available excitatory synapses in feedback projections to V1 (22% recurrent excitatory synapses in layer 2 / 3 vs a maximum of 17.2% feedback synapses; [1]). In addition, putative feedback inputs would need to be wired with high functional specificity; this degree of anatomical specificity has not been demonstrated experimentally.

### Computational role of inhibitory connectivity and physiology

Non-specific connectivity between excitatory and inhibitory neurons, as assumed in our simulation models, is consistent with the concept that inhibitory neurons simply integrate neuronal responses in the surrounding population [73], and is also consistent with experimental observations of weakly tuned or untuned inhibition in rodent visual cortex [40,50,57,64,65]. Although specific E↔I connectivity has been observed in rodent cortex [16,28], the majority of E↔I synapses are likely to be made functionally non-specifically in line with the high convergence of E→I and I→E connections observed in cortex [39,40,64].

In our models, shared inhibition is crucial to mediate competition between excitatory subnetworks (Fig. 2); inhibition is untuned because excitatory inputs to the inhibitory population are pooled across subnetworks. Poorly tuned inhibition, as expressed by the dominant class of cortical inhibitory neurons (parvalbumin expressing neurons), therefore plays an important computational role and is not merely a stabilising force in cortex.

Other inhibitory neuron classes in cortex (e.g. somatostatin or vaso-intestinal peptide expressing neurons) have been shown to exhibit feature-selective responses [57,74,75]. Recent computational work examined the influence of multiple inhibitory neuron classes with different physiological and anatomical tuning properties in a model for rodent cortex [76]. They examined the role of inhibitory connectivity on divisive and subtractive normalisation of network activity in a network with specific, orientation-tuned inhibitory connectivity. They found that specific inhibitory feedback could lead to divisive normalisation of network activity, while non-specific inhibitory feedback could lead to subtractive normalisation.

However, the computational role of specific inhibition is likely to rest on the precise rules for connectivity expressed between excitatory and inhibitory neurons. If the rules for E↔E and E↔I connections align, then a specific inhibitory population could act as a break on excitation within a subnetwork, and could allow more specific anatomical connectivity to persist while maintaining the balance between excitation and inhibition in cortex. The functional profile of this balancing pool would be highly tuned, and be similar to that of the excitatory neurons in the subnetwork, suggesting a physiological signature of specific inhibitory feedback that could be sought experimentally. Alternatively, if E↔I connection rules result in counter-tuned specificity, these connections would act to strengthen competition between subnetworks.

### Existing models of specific connectivity

As discussed above, our “like-to-like” model of orientation-tuned selective excitatory connectivity coupled with non-specific inhibitory feedback is similar in network topology to classical ring models of orientation tuning in visual cortex (e.g [51,52,77]). The principal difference in our model is the embedding of functionally selective connectivity within spatially-constrained anatomical connectivity. We showed that under model parameters chosen to be realistic in mouse V1, only a small fraction of excitatory synapses must be specific in order to introduce selective amplification and competition within the network.

Several previous models designed for columnar visual cortex have incorporated selective excitatory connectivity, either with connectivity relying on purely anatomical constraints (e.g [78]) or mimicking the spatially periodic, long-range lateral excitatory projections found in monkey, cat and other species (e.g. [79-82]). These models often incorporate assumptions of long-range inhibitory projections, which have not been described for rodent cortex, and can modify the computational properties of such models [44]. These earlier models have not examined the consequences of higher-order connectivity patterns on visual coding, and instead looked at the effect of long-range connections spanning visual space in columnar visual cortex. Our models focussed on the effect of recurrent excitatory connectivity on neurons encoding overlapping regions of visual space, and make use of the salt-and-pepper functional architecture of rodent visual cortex [14].

### Feature binding to detect higher-order visual statistics

In visual cortex of primates, carnivores and rodents, orientation tuning develops before postnatal eye opening and in the absence of visual experience [83,84]. Local recurrent connections develop after the onset of visual experience and maintain their plasticity into adulthood [83,85-89]. Statistical correlations in natural scenes might therefore lead to wiring of subnetworks under an activity-dependent mechanism such as spike-time dependent plasticity (STDP) [90-94]. Along these lines, examinations of the development of specific excitatory connections after eye opening found that similarities in feedforward input were progressively encoded in specific excitatory connections [22].

We expect that, as the specificity of lateral connections forms during development, the emergence of compound feature selectivity will gradually occur after the onset of sensory experience. This hypothesis is consistent with experiencedependent development of modulatory effects due to natural visual stimulation outside of the classical receptive field, as observed in mouse V1 [95]. A complete factorial combination of all possible features occurring in natural vision is clearly not possible. However, the most prominent statistical features of cortical activity patterns could plausibly be prioritised for embedding through recurrent excitatory connectivity. At the same time, competition induced by non-specific shared inhibition will encourage the separation of neurons into subnetworks. In our interpretation, single subnetworks would embed learned relationships between external stimulus features into functional ensembles in cortex, such that they could be recovered by the competitive mechanisms we have detailed.

In pre-frontal cortex, compound or mixed selectivity of single neurons to combinations of task-related responses has been found in several studies [96,97]. This is proposed to facilitate the efficient decoding of arbitrary decision-related variables. Binding feedforward cortical inputs into compound representations, as occurs in our “feature-binding” model, is therefore a useful computational process with general applicability. Our works suggests that specific local excitatory connectivity could be a general circuit mechanism for shaping information processing in cortical networks.

## Materials and Methods

### In-vivo calcium imaging

Experimental procedures followed institutional guidelines and were approved by the Cantonal Veterinary Office in Zürich or the UK Home Office. Procedures for urethane anaesthesia, craniotomies, bulk loading of the calcium indicator, as well as for *in vivo* two-photon calcium imaging and *in vitro* recording of synaptic connection strength were the same as described previously [24,30,98,99].

#### Preparation and imaging with OGB

Male and female three-month old wild type C7BL/6 mice were sedated with chlorprothixene (10 mg / ml in Ringer solution; 0.01 ml per 20g by weight) then anaesthetised with urethane (10% in isotonic saline; initial dose 0.1 ml per 20g by weight; supplemented as required to maintain anaesthesia). The body temperature of anaesthetised animals was monitored and controlled using a heating pad and rectal thermometer. Atropine was given to reduce secretions (0.16 ml per 20g by weight). Intrinsic optical imaging was used to locate primary visual cortex, and a craniotomy was made over V1. Briefly, the skull above the estimated location of V1 was thinned and we illuminated the cortical surface with 630 nm LED light, presented drifting gratings for 5 s, and collected reflectance images through a 4× objective with a CCD camera (Toshiba TELI CS3960DCL).

We performed bulk loading of the synthetic calcium indicator Oregon GreenBAPTA–1 (OGB–1; Invitrogen). Several acute injections of OGB–1–AM were made under visual guidance into the visual cortex [100]. Sulforhodamine (SR–101; Invitrogen) was applied topically to the pial surface, to provide labelling of the astrocytic network [101]. Time-series stacks recording activity in layer 2/3 cortical neurons were acquired at a 4–10 Hz frame rate with a custom-built microscope equipped with a 40× objective (LUMPlanFl/IR, NA 0.8; Olympus) and an 80 MHz pulsed Ti:Sapphire excitation laser (MaiTai HP; Spectra Physics, Newport). Acquisition of calcium transients was performed using custom-written software in LabView (National Instruments), and analysis was performed using the open-source FocusStack toolbox [102].

#### Preparation and imaging with GCaMP6

Adult male mice (P75–P90) were initially anaesthetized with 4–5% isoflurane in O_2_ and maintained on 1.5–2% during the surgical procedure. The primary visual cortex (V1) was localized using intrinsic imaging.

A craniotomy of 3–4 mm was opened above the region of strongest intrinsic signal response, which we assumed to be centred over V1. We then injected the genetically encoded calcium indicator GCaMP6m [103] (AAV1.Syn.GCaMP6m.WPRE.SV40; UPenn) around 250 μm below the cortical surface to target superficial layer neurons. 2–3 injections were made in a single animal and a volume of approximately 200nl was injected at each location. The craniotomy was sealed with a glass window and a metal post for head fixation was implanted on the skull with dental acrylic, contralateral to the cranial window.

For imaging, animals were anaesthetised with isoflurane at 4% for induction, then head fixed. Isoflurane concentration was lowered to 0.5–0.75% during the experiment. We maintained the animal’s body temperature at 37°C using a rectal thermometer probe and a heating pad placed under the animal. Silicon oil was applied to the eyes to keep them moist.

#### In vivo/ in vitro characterisation of function and connectivity

Methods for obtaining visual responses *in vivo* and measuring synaptic connectivity *in vitro* are described in [24]. Briefly, young C75/BL6 mice (P22–26) were anaesthetised (fentanyl, midazolam and medetomidine) and injected with OGB calcium indicators, lightly anaesthetised with isoflurane (0.3–0.5%) and head fixed. Twophoton imaging of calcium responses was used to record the response of neurons to a sequence of natural images (1800 individual images). After *in vivo* imaging experiments, the brain was rapidly removed and sliced for *in vitro* recording. Zstacks recorded *in vivo* were matched with Z–stacks recorded *in vitro* in order to locate functionally characterised neurons for electrophysiological recording. Simultaneous whole-cell recordings of up to six neurons at a time were performed. Evoked spikes and recorded EPSPs were used to identify synaptically connected pairs of neurons.

### Visual stimulation

Visual stimuli for receptive field characterisation, drifting gratings and plaids and masked natural movies were displayed on an LCD monitor (52.5 × 29.5 cm; BenQ) placed 10–11 cm from the eye of the animal and covering approximately 135 × 107 visual degrees (v.d.). The monitor was calibrated to have a linear intensity response curve. Contrast-oscillating grating and plaid stimuli were presented on an LCD monitor (15.2 × 9.1 cm; Xenarc) placed 9 cm from the eye of the animal and covering 80 × 54 v.d. The same screen was used for stimulus presentation during intrinsic imaging to locate visual cortex and during two-photon imaging. The open-source StimServer toolbox was used to generate and present visual stimuli via the Psychtoolbox package [102,104].

Stimuli for receptive field characterisation comprised a 5 × 5 array of masked high contrast drifting gratings (15 v.d. wide; overlapping by 40%; 9 v.d. per cycle; 1 Hz drift rate; 0.5 Hz rotation rate) presented for 2 s each in random order, separated by a blank screen of 2 s duration, with 50% luminance (example calcium response shown in Fig. S3a). Frames were averaged during the 2 s stimulus window to estimate the response of a neuron.

Full-field high-contrast drifting gratings (33.33 v.d. per cycle; 1 Hz drift rate) were presented drifting in one of 8 directions for 2 s each in random order, separated by a 6 s period of blank screen with 50% luminance (example calcium response shown in Fig. S3b). Frames were averaged during the 2 s stimulus window to estimate the response of a neuron.

Full-field 50% contrast drifting sine-wave gratings (25 v.d. per cycle; 1 Hz drift rate) were presented drifting in one of 16 directions for 1 s each in random order (calcium responses shown in Fig. S1). Full-field drifting plaid stimuli were constructed additively from 50% contrast sine-wave grating components (25 v.d. per cycle; 1 Hz drift rate; 1 s duration; Fig. S1). Three frames were averaged following the peak response (384 ms window) to estimate the response of a neuron.

Full-field natural movies consisted of a 43 s continuous sequence with three segments (example calcium response shown in Fig. S3c).

Full-field contrast-oscillating square-wave gratings and plaid stimuli were composed of bars of 8 v.d. width which oscillated at 2 Hz between black and white on a 50% grey background, and with a spatial frequency of 20v.d./cycle (example calcium response shown in Fig. 4a). On each subsequent oscillation cycle the bars locations shifted phase by 180°. Static gratings were used to avoid introducing a movement component into the stimulus. A base orientation for the gratings of either horizontal or vertical was chosen, and five orientations spanning ± 40deg. around the base orientation were used. Contrast-oscillating plaids were composed of every possible combination of the five oscillating grating stimuli, giving 5 grating and 10 plaid stimuli for each experiment. A single trial consisted of a blank period (50% luminance screen) presented for 20s, as well as presentations of each of the gratings and plaids for 5 s each, preceded by 5 s of a blank 50% luminance screen, all presented in random order. Frames from 0.25 s to 4.75 s during the stimulus period were averaged to estimate the response of a neuron.

### Analysis of calcium transients

Analysis of two-photon calcium imaging data was conducted in Matlab using the open-source FocusStack toolbox [102]. During acquisition, individual twophoton imaging trials were visually inspected for Z-axis shifts of the focal plane. Affected trials were discarded, and the focal plane was manually shifted to align with previous trials before acquisition continued. Frames recorded from a single region were composed into stacks, and spatially registered with the first frame in the stack to correct lateral shifts caused by movement of the animal. Only pixels for which data was available for every frame in the stack were included for analysis. A background fluorescence region was selected in the imaged area, such as the interior of a blood vessel, and the spatial average of this region was subtracted from each frame in the stack. The baseline fluorescence distribution for each pixel was estimated by finding the mean and standard deviation of pixel values during the 10 s blank periods, separately for each trial. Regions of interest (ROIs) were selected either manually, or by performing low-pass filtering of the OGB (green) and sulforhodamine (red) channels, subtracting red from green and finding the local peaks of the resulting image.

A general threshold for responsivity was computed to ensure that ROIs considered responsive were not simply due to neuropil activity. The responses of all pixels outside any ROI were collected (defined as “neuropil”), and the Z-scores of the mean Δ*F/F*_0_ responses during single visual stimulus presentations were computed per pixel, against the baseline period. A threshold for single-trial responses to be deemed significant (*z*_trial_) was set by finding the Z-score which would include only 1% of neuropil responses (*α* = 1%). A similar threshold was set for comparison against the strongest response of an ROI, averaged over all trials (*z*_max_). These thresholds always exceeded 3, implying that single-trial responses included for further analysis were at least 3 standard deviations higher than the neuropil response. Note that this approach does not attempt to subtract neuropil activity, but ensures that any ROI used for analysis responds to visual stimuli with calcium transients that can not be explained by neuropil contamination alone.

The response of AN ROI to a stimulus was found on a trial-by-trial basis by first computing the spatial average of the pixels in AN ROI for each frame. The mean of the frames during the blank period preceding each trial was subtracted and used to normalise responses (Δ*F/F*_0_), and the mean Δ*F/F*_0_ of the frames during the analysed trial period was computed. The standard deviation for the baseline of a neuron was estimated over all Δ*F/F*_0_ frames from the long baseline period and the pre-trial blank periods. ROIs were included for further analysis if the ROI was visually responsive according to trial Z-scores (maximum response > *z*_max_) and reliable (trial response > *z*_trial_ for more than half of the trials). The response of a neuron to a stimulus was taken as the average of all single-trial Δ*F/F*_0_ responses.

Receptive fields of neurons recorded under natural movie and drifting grating stimulation were characterised by presenting small, masked high-contrast drifting gratings from a 5 × 5 array, in random order (see above; Fig. S3a). A receptive field for each neuron was estimated by a Gaussian mixture model, composed of circularly symmetric Gaussian fields (*ρ* = 7.5 v.d.) placed at each stimulus location and weighted by the response of the neuron to the grating stimulus at that location. The centre of the receptive field was taken as the peak of the final Gaussian mixture. Neurons were included for further analysis if the centre of their receptive field lay within a 7.5 v.d. circle placed at the centre of the natural movie visual stimulus. Example single-trial and trial-averaged calcium responses to natural movie stimuli are shown in Fig. S3c.

### Response similarity measures and response metrics

The similarity in response between two neurons was measured independently for grating and plaid stimuli. The set of grating responses for each neuron were composed into vectors *R*1_*g*_ and *R* 2_*g*_, where each element of a vector was the trial-averaged response of a neuron to a single grating orientation. The similarity in grating responses between two neurons was then given by the Pearson’s correlation coefficient between *R*1_*g*_ and *R*2_*g*_: *ρ*_*g*_ =corr(*R*1_*g*_, *R*2_*g*_) (see Fig. S3b, inset). The similarity in response to plaid stimuli was computed analogously over the sets of trial-averaged plaid responses *R*1_*p*_ and *R* 2_*p*_: *ρ*_*p*_=corr(*R*1_*p*_, *R* 2_*p*_) (see Fig. 4a, inset). Similarity was only measured between neurons recorded in the same imaging site.

The similarity between neurons in their responses to movie stimuli (*ρ*_m_) was measured by computing the signal correlation as follows. The calcium response traces for a pair of neurons were averaged over trials. The initial 1 s segment of the traces following the onset of a movie segment were excluded from analysis, to reduce the effect of transient signals in response to visual stimulus onset on analysed responses. The Pearson’s correlation coefficient was then calculated between the resulting pair of traces (*ρ*_m_; see Fig. S3c, inset). Note that correlations introduced through neuropil contamination were not corrected for, with the result that the mean signal correlation is positive rather than zero. For this reason we used thresholds for “high” correlations based on percentiles of the correlation distribution, rather than an absolute correlation value.

The similarity between neurons in their responses to flashed natural stimuli (*ρ*_Ca_; Fig. S3f) was measured as the linear correlation between the vector of responses of a single neuron to a set of 1800 natural stimuli [24].

The Orientation Selectivity Index (OSI) of a neuron was estimated using the formula *OSI* =[max(*R*_*g*_)-min(*R*_*g*_)] sum(*R*_*g*_), where *R*_*g*_ is the set of responses of a single neuron to the set of grating stimuli. The OSI of a neuron ranges from 0 to 1, where a value of 1 indicates that a neuron responds only to a single grating stimulus; a value of 0 indicates equal, nonselective responses to all grating stimuli.

The Plaid Selectivity Index (PSI) of a neuron, describing how selective a neuron is over a set of plaid stimuli, was calculated using the formula *PSI*=1-[-1+∑ *R*_*p*, *j*_ max(*R*_*p*_)] [ *#*(*R*_*p*_)–1], where *#*(*R*_*p*_) is the number of stimuli in *R*_*p*_ [30]. The PSI of a neuron ranges from 0 to 1, where a value of1 indicates a highly selective response, where a neuron responds to only a single plaid stimulus; a value of 0 indicates equal, nonselective responses to all plaid stimuli.

A plaid Modulation Index (MI), describing the degree of facilitation or suppression of a neuron in response to plaid stimuli, was calculated using the formula *MI* =[ max(*R*_*p*_)-max(*R*_*g*_)] [ max(*R*_*p*_)+max(*R*_*g*_)], where *R*_*p*_ is the set of responses of a single neuron to the set of plaid stimuli [30]. The MI of a neuron ranges from –1 to 1. Values of MI < 0 indicate stronger responses to grating stimuli compared with plaid stimuli; values of MI > 0 indicate stronger responses to plaid stimuli. A value of MI = –1 indicates that a neuron responds only to grating stimuli; a value of MI = 1 indicates that a neuron responds only to plaid stimuli.

The proportion of facilitating and suppressing neurons was compared between mouse V1 and model responses using two-tailed Fisher’s exact tests. The population of responsive neurons was divided into three groups: facilitating (MI > 0.05); suppressing (MI < –0.05); and non-modulated (–0.05 <= MI <= 0.05). These categories were arranged into three 2 × 3 contingency tables, with each table tested to compare facilitation and suppression between mouse V1 and one model.

### Generation of V1 control responses

We used single-cell, single-trial responses to oscillating contrast grating stimuli to explore whether we could distinguish between correlated and decorrelated responses to plaid stimuli, given experimental variability and noise. For each cell in the experimentally-recorded data set, we used the set of grating responses R_g_ to generate plaid responses R_p_ for the same cell, under the assumption that the response to a plaid was linearly related to the sum of the responses to the two grating components. For each plaid, we randomly selected a single-trial response for each of the grating components of the plaid. The predicted single-trial plaid response was the sum of the two grating responses. We generated 100 bootstrap samples for each experimental population, with each sample consisting of the same number of trials and neurons as the experimental population. We then quantified the relationship between grating and plaid responses as described for the experimental data.

### Statistical methods

We used a sample size commensurate with those used in the field, and sufficient for statistical analysis of our observations. No explicit sample size computation was performed.

Two-sided, non-parametric statistical tests were used unless stated otherwise in the text.

## Acknowledgements

The authors gratefully acknowledge T Mrsic-Flogel, L Cossell and MF Iacaruso for providing the data analysed in Fig. S3f,g, and gratefully acknowledge WC L Lee and C Reid for providing the data analyzed in Fig. S3h. We are grateful to MA Penny and the attendees of the CapoCaccia workshop for helpful discussions.

---

**Supplementary Figure 1:**
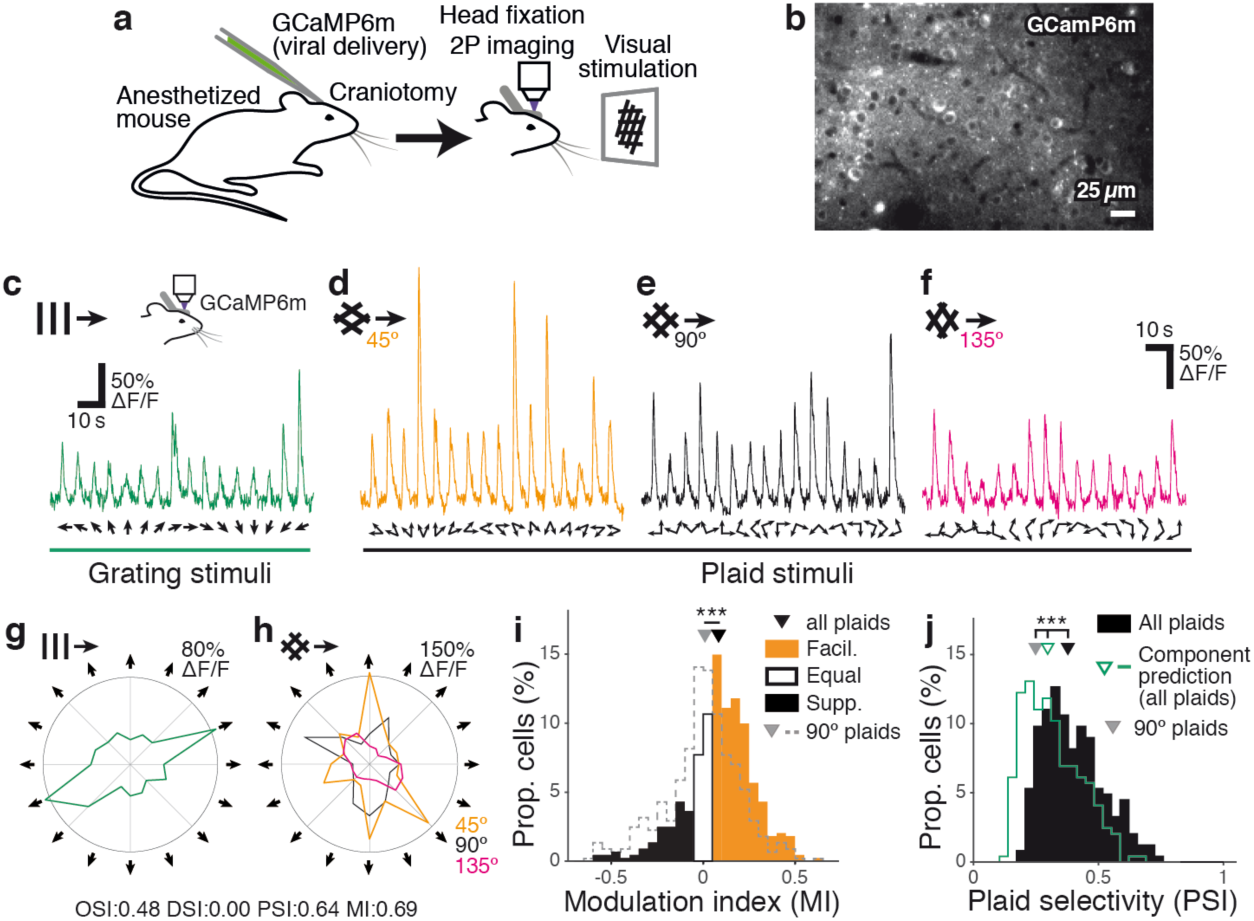
Plaid responses are facilitatory and selective in mouse VI. **(a, b)** We re-corded responses from layer 2/3 neurons in Vl using two-photon imaging of animals with viral delivery of GCaMP6m (8 animals, 8 sessions, 441 / 879 responsive *I* imaged neurons; see Methods.) **(c-h)** We probed mouse Vl with grating component stimuli composed of grating stimuli with 16 drift directions, and three full sets of plaid stimuli composed of 45°, 90° and 135° relative grating component orientations. We defined a modulation index (MI) to quantify the degree of facilitation or suppression elicited by plaid stimuli over grating stimuli, for single cortical neurons; large positive values for MI indicate strong facilitation in response to plaid stimuli, whereas large negative values indicate strong suppression (see Methods). Visual responses to the full set of plaid stimuli were dominated by facilitation, and were signif icantly more facilitatory than when considering only the set of 90° plaids (median modulation index MI 0.098± [0.081 0.12] vs 0.011± [-0.0060 0.027]; p < 1X10-10, Wilcoxon rank-sum; all following values are reported as median± 95% bootstrap confidence intervals unless stated otherwise). (**j**) The presence of stronger facilitation when comparing responses to the full set of plaid stimuli with responses to 90° plaids alone is consistent with our earlier finding that some neurons in mouse Vl are highly selective for particular combinations of grating components [31]. Accordingly, we used a plaid selectivity index (PSI) to quantify how selective were the responses of single neurons over the set of plaid stimuli (see Methods). The PSI was defined in analogy to orientation or direction selectivity indices (OSI or DSI), such that val-ues of PSI close to 1 indicate that a neuron responds to only a single plaid stimulus out of the set of pre-sented plaid stimuli. Values of PSI close to O indicate that a neuron responds equally to all plaid stimuli. Responses to the full set of plaid stimuli were highly selective; significantly more selective than predicted by a component model generated using all plaid and grating stimuli [90] (median PSI 0.38± [0.36 0.41] vs 0.30± [0.28 0.31]; p < 1X10-10, Wilcoxon rank-sum) and indeed significantly more selective than responses to the 90° plaids alone (median 90° PSI 0.25± [0.23 0.28]; p < 1X10-10 vs all plaids, Wilcoxon rank-sum).*** p < 1X10-10. OSI: orientation selectivity index; DSI: direction selectivity index; PSI: plaid selectivity index; MI: modulation index; facil.: facilitating; supp.: suppressing; prop.: proportion.

**Supplementary Figure 3:**
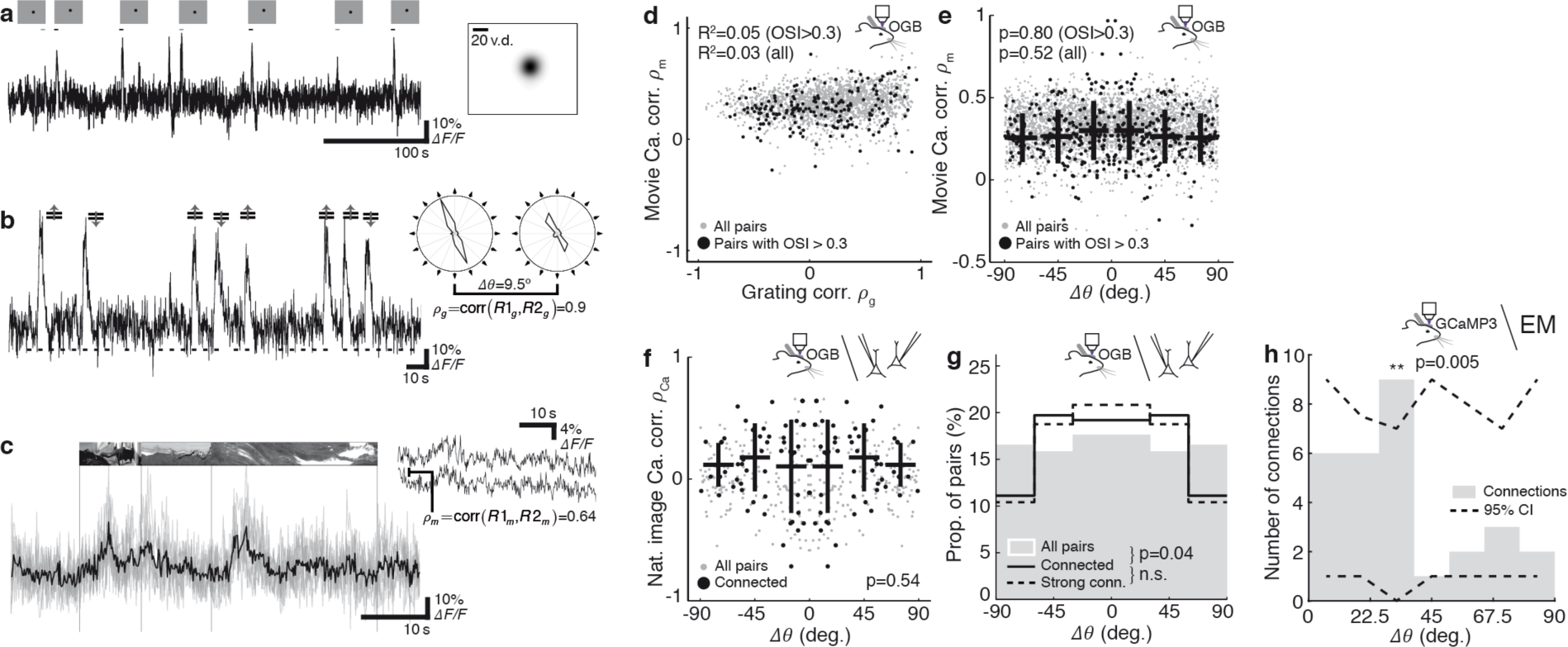
Estimated parameters for cortex place it in an Inhibition-Stabilised Network (ISN) regime, with competition provided by specific excitatory connectivity. **a** The network stability regimes in the parameter space defined by total inhibitory weight *g*_1_·*n*_1_ and total excitatory weight *g*_E_·*n*_E_ for a random network (proportion of specific synapsess= 0%). Nominal parameter estimates for rodent cortex (cross) place the network in a regime that requires inhibitory feedback for stability (an ISN; [39]), but which does not lead to competition between excitatory neurons. Inhibition must be unrealistically strengthened to obtain competition (100x and 200x estimates for rodent cortex; top of panel; shading indicates competition). However, over–ly-strong inhibition leads to inhibition-driven oscillations (IO). **b** When the proportion of specific synapsessis raised to 20%, nominal parameters for rodent cortex permit competition (shading indicates strength of competition). Note that the maximum excitatory strength permitted while maintaining network stability is reduced. **c** When *s* = 40%, nominal parameters for rodent cortex become unstable (cross is just inside unstable region). **d** Network stability regimes for the parameter space defined bysand *g*_E_·*n*_E_, with nominal value chosen for *g*_1_ ·*n*_1_ (crosses in a-c). Nominal value for *g*_E_·*n*_E_ is indicated by a dashed line. Both excitatory strength *g*_E_·*n*_E_ and the proportion of specific synapses s affect network stability and the strength of competition. Abbreviations: *g*_I_.E: Synaptic strength per inhibitory or excitatory synapse; *n*_1.E_: Number of synapses made by each inhibitory or excitatory neuron; AS: Intrinsically stable network, stable in the absence of inhibition;

**Supplementary Figure 2:**
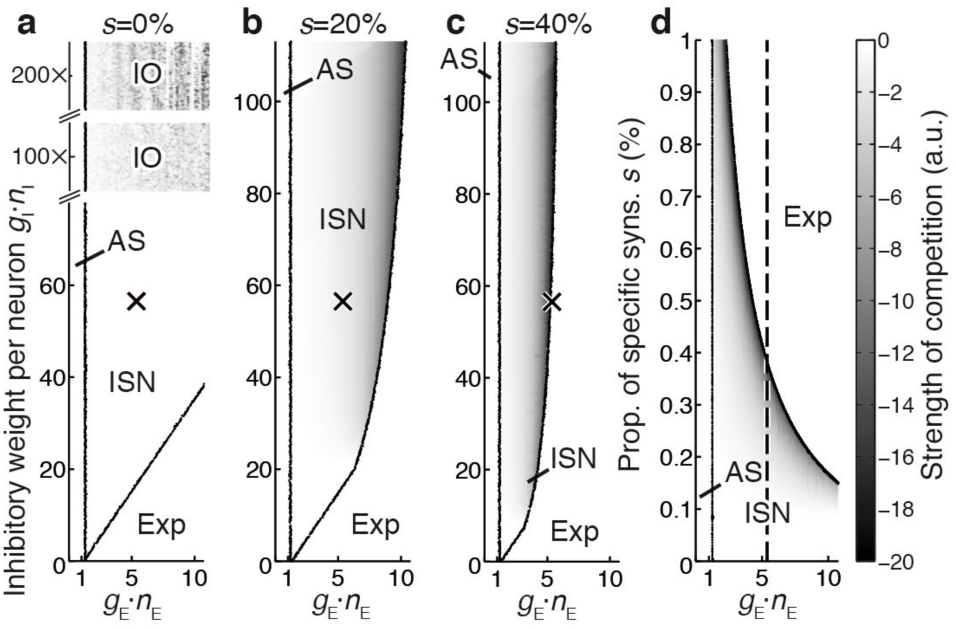
Connected neurons span a wide range of preferred orientations in mouse VI. **a** Characterisation of receptive field location using sparse drifting/rotating grating stimuli. Single-trial 0GB calcium responses (black); presentation time of optimal stimulus and sub-optimal stimulus indicated (black and grey bars). Right inset: estimated RF location for the same neuron. **b** Single-trial 0GB calcium response to drifting grating stimuli (black); presentation of optimal stimulus orientation indicated above, all stimulus presentation times indicated below. Right inset: calculation of grating response similarity *ρ* between two neurons. **c** Single-trial (grey) and trial-averaged 0GB calcium response (black) to natural movie stimuli. Vertical lines indicate timing of movie sequence onset. Right inset: calculation of movie response similarity (pm), using signal correlations over trial-averaged responses from two neurons. **d** Pairs of neurons with high signal correlations to natural movies (pm), which predicts a high probability of connection [21], can have similar or dissimilar grating responses. Pairs of neurons with similar orientation preference are not more likely to have high Pm **(e)** or high signal correlation to flashed natural scenes Pea (**f**) than pairs with dissimilar orientation preference. **g** Connected pairs are slightly more likely to share similar orientation preferences than unconnected pairs [21,24], but nevertheless span almost arbitrary orientation differences (:::c:20% of pairs with close to orthogonal orientation preference). **h** In data from functionally characterized neurons with connections reconstructed under electron microscopy[105], connected pairs are more likely to share similar preferred orientations. An excess of connections was present at orientation preference differences of around 30_°_ (p = 0.005, Monte-Carlo test). Dashed lines: 95% boostrap confidence intervals (CI). d-e: in vivo two-photon calcium imaging; f-g: in vivo calcium imaging coupled with in vitro simultaneous patching to detect connected pairs; data from [24]. h: in vivo calcium imaging coupled with electron microscopy (EM) reconstruction to identify conencted neurons; data from [105]. e-f: Kruskal-Wallis tests; g: Ansari-Bradley test; h: Monte-carlo test. n.s.: p > 0.05. Strong connections: strongest 50% of connected pairs, measured by EPSP amplitude. Corr: correlation; conn.: connection.

**Supplementary Figure 4:**
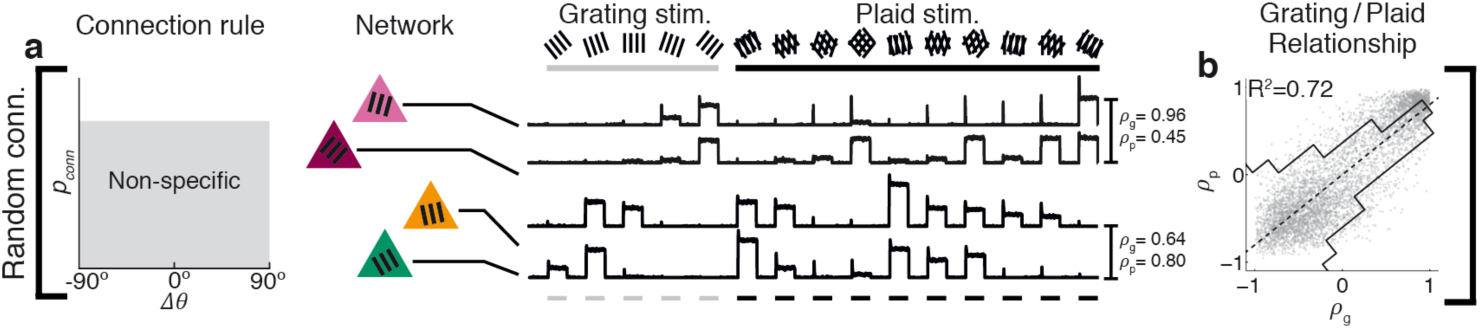
Grating and plaid responses are highly correlated in a model with random connectivity. **a** Under the non-specific connectivity model, synapses between pairs of neurons are formed without regard to functional response similarity of the neurons. Neurons form synapses stochastically, according to spatial proximity. Two example pairs of neurons are shown, and their responses to a set of grating and plaid stimuli. **b** Neurons with similar responses to grating stimuli (high *ρ*_g_) have similar responses to plaid stimuli (high *ρ*_p_), and vice versa.

